# EOLA1 functions in nucleotide salvage through deacetylating free N4-acetylcytidine

**DOI:** 10.64898/2026.08.01.741249

**Authors:** Sebastien Relier, Sarah Schiffers, Hamid Beiki, Maria Prigge, Nishu Tyagi, Cyrinne Achour, Ayush Raman, A. Maxwell Burroughs, L. Aravind, Shalini Oberdoerffer

## Abstract

RNA-based medicines rely on modified nucleotides to promote immune evasion and in vivo efficacy. Nucleotides generated from RNA degradation are either exported or recycled through metabolically favorable salvage pathways, though whether modified nucleotides are efficiently recycled remains unclear. N^4^-acetylcytidine (ac⁴C) is a naturally occurring modification in rRNA and tRNA that has shown promise in therapeutic mRNA applications. However, N^4^-acetylation impairs cytidine deamination, the first step in cytidine salvage. Here, we investigate the endogenous mechanisms that enable ac⁴C metabolism. Through sensitive sequence and structural analyses, we identify the uncharacterized human ASCH domain protein EOLA1 as a key ac⁴C deacetylase in nucleotide salvage. EOLA1 inactivation leads to free intracellular ac⁴C accumulation and increased cytotoxicity upon nucleotide export inhibition. While steady-state ac⁴C levels in cellular RNAs remain unchanged, EOLA1-dependent regulation of free ac⁴C is evident basally and is exacerbated by exogenous mRNA delivery. Proteomic analyses place EOLA1 in proximity to ribosomal proteins, adjacent to endogenous ac⁴C sources. In vitro assays confirm EOLA1 specificity for ac⁴C, and structural analysis reveals a narrow nucleotide-binding pocket consistent with mononucleotide selectivity. These findings identify EOLA1 as a bona fide ac⁴C eraser and uncover a previously unrecognized pathway for recycling modified nucleotides with relevance to therapeutic RNA design.

## Introduction

Cells continuously turn over their RNA, releasing free ribonucleotides that are either exported or recycled through nucleotide salvage pathways (1,2). Because de novo nucleotide synthesis is energetically costly, salvage predominates in most somatic tissues, and ribosomal RNA degradation is a major source of the recycled pool (1). RNA, however, carries a diverse array of chemical modifications, and whether modified nucleotides can be efficiently routed through salvage, or must instead be excreted to prevent their misincorporation into nascent nucleic acids, remains poorly understood (2,3). This question is gaining practical significance, as RNA-based medicines depend on modified nucleotides to evade innate immune recognition and sustain translation in vivo (4,5). The intracellular fate of the modified nucleotide byproducts released upon turnover of such therapeutic RNAs is therefore directly relevant to their design.

N4-acetylcytidine (ac⁴C) is a universally conserved RNA modification and the sole known acetylation event in eukaryotic RNA (6). It is installed by NAT10 (Kre33 in yeast, TmcA in bacteria) across tRNA, rRNA, and human mRNA (7–12), where it stabilizes Watson:Crick pairing with guanosine to promote tRNA decoding and modulate mRNA translation (10,11,13,14). This same stabilizing property has recently made ac⁴C an attractive component of synthetic mRNA: we demonstrated that ac⁴C-modified mRNA suppresses innate immune activation as effectively as the industry-standard N1-methylpseudouridine (m¹Ψ) while supporting higher and more faithful translation (15). ac⁴C thus represents both an endogenous mark and an emerging therapeutic building block, raising the question of how acetylated cytidine is metabolized once liberated from RNA.

The first and committed step of cytidine salvage is the deamination of cytidine to uridine by cytidine deaminase (16). N4-acetylation occludes the exocyclic amine targeted by this reaction, rendering acetylated cytidine refractory to deamination (17). Consequently, ac⁴C released during RNA turnover cannot directly re-enter salvage and must be either exported or enzymatically deacetylated before it can be recycled. Whether human cells possess a dedicated activity for this purpose, however, is unknown. A precedent exists in bacteria: in a screen of acylated derivatives, the *E. coli* protein YqfB was found to preferentially deacetylate free ac⁴C in vitro (18). YqfB is composed entirely of a standalone "activating signal cointegrator-1 homology" (ASCH) domain, a pseudouridine synthase (PUA)-like β-barrel fold predicted to bind modified nucleobases in nucleic acids and nucleotides (18–23). Although the ASCH domain is conserved across all domains of life and humans encode multiple ASCH-containing proteins, sequence searches have not identified a direct human ortholog of YqfB (19,24,25).

Here, we identify human EOLA1 as a first-in-class nucleoside deacetylase belonging to a distinct clade of ASCH domains from the bacterial YqfB. In vitro analyses with purified protein confirmed EOLA1 specificity for ac⁴C, while cellular studies involving EOLA1 overexpression or ablation established its endogenous function in deacetylating free ac⁴C derived from RNA degradation. Notably, EOLA1 does not measurably erase ac⁴C from intact cellular RNAs, consistent with a role restricted to free nucleotides rather than RNA-incorporated modifications. Our findings add ac⁴C to the growing list of modified nucleosides subject to dynamic metabolic regulation and uncover the potential of nucleotide deacetylases in modified-nucleotide recycling and in mitigating the toxicity of RNA-based therapeutics.

## Material and Methods

### Comparative genomics and domain identification

Domains were identified using a collection of HMMs and PSSMs maintained by the Aravind lab, along with HMMs from the Pfam database, by means of the RPSBLAST and HMMSCAN programs. To further refine detection, domain identification was extended through remote homology analysis using HHpred against profiles built from the Pfam and PDB databases. Genomic neighborhoods were extracted from genomes deposited in the NCBI Genome database using custom Python scripts. Further analysis of these genomic neighborhoods was performed by clustering the protein products of neighboring genes.

### Domain architectures and phylogenetic analysis

Phylogenetic analysis of the ASCH domains was performed using the FastTree program and rendered using TreeViewer. The domain architectures and conserved gene neighborhoods were compiled from the significant matches in the above-described profile searches and superimposed onto branches of the trees. The branches were colored according to their prevalent contextual connections and rooted based on the conservation pattern of the active motif of the ASCH domain.

### Pairwise alignment of protein structures

Human PDB structures were downloaded from RCSB protein data bank (26) or AlphaFold (27). The structure of YqfB (PDB : 1te7, AlphaFold : Q8TE69-model-v1) was aligned to any human structures from RCSB and AlphaFold using MMLigner version 1.0.2 (28). The pairwise alignment with the highest compression score is considered the most significant. Structure alignment of favorite targets was further visualized using Pymol 3.1.

### Plasmid Construction

The coding sequences of EOLA1 and EOLA2 were cloned into pCDNA5 FRT/TO through HindIII and BamHI digestion and ligation, followed by transformation in DH5α and selection for ampicillin resistance. Plasmid DNA was isolated by MiniPrep (Zymo Research) and validated through Sanger sequencing (Psomagen). EOLA1 was further mutated using the QuickChange II mutagenesis kit (Agilent) to generate a plasmid encoding a R26A catalytic mutant (Table S1).

### Cell culture and transfection

HeLa were maintained in DMEM supplemented with 10% bovine calf and 50 uM glutamine. For si-RNA transfections, 150,000 cells were seeded in a 6-well plate and transfected the same day with 25 nM siRNA targeting EOLA1 (5’- UACAUUAGGUUAAAUCUGAtt-3’) using lipofectamine RNAimax (ThermoFisher Scientific). Cells were assayed after a 72h incubation. For plasmid DNA transfections, 150,000 cells were seeded in a 6-well and transfected 2 days later with 2ug of plasmid DNA using lipofectamine 2000 (ThermoFisher Scientific). Cells were assayed after a 24h incubation.

### Monocyte isolation

Primary human monocytes were isolated from peripheral blood of anonymous donorsutilizing the EasySep™ Direct Human Monocyte Isolation Kit (Stem Cell Technologies, Cat. #: 19669) according to manufacturer’s instructions, followed by addition of 25 mM HEPES (Quality Biological, Cat. #: 118-089-721), 50 ng/mL recombinant human IL4 (Peprotech, Cat. #:200-04) and 50 ng/mL recombinant human GM-CSF (Sigma-Aldrich, Cat. #: GF-304) in complete RPMI to yield monocyte-derived dendritic cells.

### Flow Cytometry

The following antibodies were used in flow cytometry: PE-conjugated anti-Hu/NHP CD25 antibody (CD25-4E3, eBioscience, Cat. #: 12-0257-42), APC-conjugated anti CD14 antibody (61D3, eBioscience, Cat. #: 17-0149-42), PerCP/Cyanine5.5-conjugated anti-human CD11c antibody (3.9, Biolegend, Cat. #: 301624), PE-conjugated anti-CD209 (DC-SIGN) antibody (9E9A8, Biolegend, Cat. #: 330105). 0.1-1x10^6^ cells were stained in 100 µL staining buffer (1%FBS in PBS) for 30 mins at room temperature in the dark, washed twice with PBS and resuspended in 1% paraformaldehyde containing staining buffer. Flow cytometry acquisition was performed on BD FACSymphony A5 and examined using FACSDiva software (BD Bioscience), data was further analyzed using FlowJo v10.8.1 (BD).

### Cell proliferation

150,000 cells were seeded in a 6-well plate and cultured for 24h, 48h,72h at 37°C/5% CO2. At each time point, cells were trypsinized and the number of viable cells was determined using Trypan Blue (Sigma Aldrich) and an automatic cell counter (Nexcelom Bioscience).

### Cell viability assay

2,000 cells were seeded in a 96-well plate and incubated for 24h at 37°C/5% CO_2_. Decreasing amount of Dipyridamole (125 to 0.25 uM), hydrogen peroxide (250 to 0.25 uM), or sodium arsenite (100 to 0.19 nM) were added to the cells, followed by an additional 72h (Dipyridamole) at 37°C/5% CO_2_. 20 uL of Cell Titer 96 (Promega) was added to the cells and incubated for 2h. Absorbance at 490 nm was measured and the IC50 was determined graphically.

### Replicative stress analysis by Immunofluorescence

100,000 cells were seeded into a 6-well plate containing a coverslip and incubated for 72h at 37°C/5% CO_2_. Cells were fixed using 4% formaldehyde for 10 min at room temperature. Cells were washed in 1X PBS, permeabilized in 1X PBS 0.25% Triton for 10min, then blocked in 1X PBS 3% BSA and 0.1% Tween for 45min. Immunostaining was performed overnight at 4°C with an anti p-ɣH2AX (Abcam, EP854(2)Y, 1;250). Cells were washed three times in 1X PBS + 0.1% tween-20 then incubated with a donkey anti-Rabbit antibody (Thermo Fisher Scientific, A10042, 1:200) for 1h at room temperature.

Each coverslip (VWR® Micro Cover Glasses, Square, No. 2, 48368-062) was mounted in 10uL anti-fade mounting medium with DAPI (Vectashield, H-1200-10). Images were acquired using an LSM780 confocal microscope (Zeiss).

### Generation of EOLA1/2 Knock-out cells

CRISPR-Cas9 mediated knock-out of the EOLA1 gene was achieved using the PX458 plasmid (Addgene) encoding pSpCas9–2A-GFP, along with a guide RNA (gRNA) targeting EOLA1 exon 4 (5‘-CACCGTCCCCTGCTGAGCAGCCAG-3’) selected through http://crispr.mit.edu. HeLa were transfected with the plasmid DNA using Lipofectamine 2000 (ThermoFisher Scientific) and GFP positive cells were sorted and collected 24h later with the FACSAriaII cell sorter (BD Biosciences). Single clones were seeded into wells of a 96-well plate in complete DMEM. Positive clones were validated by RT-qPCR, Western blotting and DNA sequencing.

### Polysome Fractionation

2 plates (150 cm2) were seeded with 2 x 10^6^ cells. After 48h, cells were treated with 20 µg/ml emetine for 5 min at 37°C, washed twice with ice-cold PBS, and scraped in ice cold PBS. Cells were centrifuged at 400g for 5 min at 4°C. Cell pellets were resuspended into 1mL of polysome lysis buffer (5% sucrose, 50mM Tris-HCl pH 7.5, 5 mM MgCl2, 25 mM KCl) supplemented with RNase-inhibitor (3U/uL) and protease-inhibitor (1X). Lysates were left on ice for 10 min before centrifugation at 13,200 rpm for 10 min at 4°C. Supernatants were loaded on 15-50% sucrose gradients and centrifuged for 2h26 at 41,000 rpm at 4°C in a SW41 rotor (Beckman Coulter). Absorbance at 260 nm was recorded using a BioComp Fractionation System and a Triax Flow Cell model FC-1 for 260 nm scans.

### *In Vitro* mRNA synthesis

#### *FLuc* RNA synthesis

300 ng of *FLuc* DNA was transcribed using the T7 Megascript kit (Thermo Fisher Scientific). Transcription reactions were incubated for 2h at 37°C and plasmid DNA was further removed with 2U of Turbo DNAse for 30 min at 37°C. RNA was precipitated using Lithium Chloride overnight at -20°C.

#### *NanoLuc* RNA synthesis

1 ug of *NanoLuc* DNA was transcribed in the presence of 5mM dNTPs, 4 mM of CleanCap AG (Trilink), 10mM DTT, 20U of Murine RNase inhibitor, 0.4U of Inorganic PyroPhosphatase and 100U of T7 RNA polymerase (NEB). Transcription reactions were carried out at 37°C for 2h. After transcription, the DNA template was digested for 30 min at 37°C in presence of 4U Turbo DNase I. RNA was further purified using RNA Clean XP Beads according to manufacturer’s instructions (Beckman Coulter).

#### Non-structured and structured RNA synthesis

DNA templates for non-structured and structured RNAs were generated by hybridization with T7 promoter at 94°C for 2min followed by a cool down to room temperature. 1ug of DNA template was transcribed in the presence of 5mM dNTPs, 4 mM of CleanCap AG (Trilink), 10mM DTT, 20U of Murine RNase inhibitor, 0.4U of Inorganic PyroPhosphatase and 100U of T7 RNA polymerase (NEB). Transcription reactions were carried out at 37°C overnight. After transcription, the DNA template was digested for 30 min at 37°C in presence of 4U Turbo DNase I. Small RNA was further purified using RNA Clean and Concentrator (Zymo Research) according to manufacturer’s instructions.

### Immunoprecipitation

HeLa were lysed in low salt buffer (50 mM Tris-HCl, 150 mM NaCl, 1 mM EDTA, 0.5% Triton-X100, pH 7.4). 500 mg HeLa lysate was incubated with 1 ug FLAG antibody conjugated to magnetic protein G beads for 18h at 4°C. Antibody-protein complexes were washed 4 times with wash buffer (50 mM Tris-HCl, 150 mM NaCl, 1 mM EDTA, 0.2% Triton-X100, pH 7.4) and further eluted with 500 uM of FLAG peptide for 2h at 4°C. Eluates were separated on 4-12 % Bis-Tris acrylamide followed by Western blotting.

### *In vitro* deacetylation

#### Synthetic RNA

250 ng *FLuc* RNA was treated with 10% of the EOLA1-FLAG IP lysates in the presence of 50 mM potassium phosphate buffer pH 8.0 and incubated for 1h at 37°C at 350 rpm. Treatment with lysates from IP control were used as negative controls.

#### Free Nucleosides

1 mM of modified nucleosides (ac^4^C, m^6^A, m^5^C) were treated with 10% of EOLA1-FLAG IP lysates in the presence of 50 mM potassium phosphate buffer pH 8.0.

Reactions were mixed with 1mM guanosine and 1mM uridine for further data normalization. Samples were incubated for 1h at 37°C at 350 rpm prior to mass-spectrometry analysis. FLAG-IP lysates were used as a negative control of spontaneous deacetylation.

### Dot-blot assay

Synthetic RNA (5025-12 ng) or total RNA (1-2 ug) was loaded onto nylon membrane (Amersham) and crosslinked twice at 120 mJ. As a loading control, the RNA was first stained with 0.1% methylene blue for 5 min at room temperature (RT). After three washes in 0.1% TBST, a colorimetric picture of the membrane was taken with a chemidoc imager.

Membranes were blocked with 0.1% TBST/5% milk for 30-60 min at RT. Immunoblotting was performed with anti-ac^4^C (1:1000), anti-m^6^A (1:1000), and anti-m^5^C (1:1000) antibodies for 16- 24h at 4°C. After three washes of 10 minutes each, membranes were incubated overnight at 4°C with anti-rabbit (1:10,000) or anti-mouse (1:10,000). Membranes were washed 3x with 0.1% TBST for 10 min each. ac^4^C signal was revealed with ECLPrime (Amersham) or ECL femto (ThermoFisher Scientific) and chemiluminescence signal was acquired using a ChemiDoc imager.

### RNA extraction

Cell pellets were resuspended in TRIzol. Chloroform (1:5) was added and samples were incubated for 5 min at RT. The upper phase (RNA) was further precipitated with 100 % ethanol, sodium acetate and linear acrylamide for 2-16h at -20°C. RNA was centrifuged at 13,200 rpm for 15 min at 4°C. Pellets were washed twice in 75% EtOH and dissolved in water.

RNA quantity and quality were assessed with Nanodrop spectrophotometer or Agilent bioanalyzer. Small RNA was enriched using the miRvana kit according to manufacturer’s instructions (ThermoFisher Scientific). mRNA was enriched using two rounds of polyA purification with oligodT beads according to manufacturer’s instructions (Invitrogen)

### Immuno-northern blot

2-10 ug of total RNA was resolved on 1% agarose in 1X denaturing Buffer (Thermo Fisher Scientific), followed by transfer to Hybond+ membrane (Amersham) for ≥ 3h at RT in the presence of 20X SSC buffer. Membranes were UV crosslinked twice at 120mJ/cm^2^ using a UV Stratalinker (Stratagene) and blocked for 30 min at RT in 5% milk/0.1% TBST. Immunostaining was performed overnight with anti-ac^4^C antibody (Abcam, 1:1,000).

Membranes were next washed three times in 0.1% TBST for 10 min, followed by incubation with anti-Rabbit-HRP antibody for ≥6h at 4° (Cell Signaling, 1:10,000). ac^4^C signal was revealed using ECL substrate (Thermo Scientific) after an additional three 10 min washes in 0.1% TBST and imaged using a ChemiDoc Imaging System (Biorad).

### RT-qPCR

1 ug of RNA was reverse transcribed with 0.5U superscript IV and 0.5 uM random hexamers. Samples were incubated at 25°C for 10 min, 50°C for 50 min and 75°C for 15 min. cDNA was diluted 20-fold and 1 uL of cDNA was used to carry out real time PCR using SYBR Green Mix (Roche). Relative quantification was achieved using the ΔΔCt method with RPS16 as a housekeeping gene (Table S1).

### RetraC:T-seq

RetraC:T-seq was performed as previously described (29). Briefly, 1-3ug of total RNA was treated for 20 min at 20°C with 100 mM NaCNBH3 (Sigma Aldrich) dissolved in 100 mM HCl. After ethanol precipitation, the treated RNA was reverse-transcribed using 0.2U of HIV reverse-transcriptase, random hexamers, and dNTPs mix composed of 500 uM ATP, 500 uM CTP, 500 uM TTP, 125 uM GTP and 375 uM 2-aminodATP (ApexBio). 25% of cDNA reaction was used to amplify 18S sequence using Q5 Hot Start PCR (NEB) and primers flanking helices 34 and 45. Amplicons were purified using PCR purification kit (Qiagen) and analyzed through Sanger sequencing (Psomagen). C:T mismatch rates at ac^4^C sites were calculated as follows: C:T = Peak Height of T / (Peak Height of C + Peak Height of T) * 100.

### Protein extraction

Cells were washed once then scraped in ice-cold PBS 1X. Cell suspensions were centrifuged for 5 min at 400g at 4°C. Cell pellets were resuspended in 50 – 100 uL of 1X RIPA buffer then left on ice for 10 min. Protein lysates were cleared by centrifugation at 16,100g for 10 min at 4°C and protein concentration quantified using bicinchoninc acid assay (Thermo FisherScientific).

### Western blot

5-50 ug of protein lysate was loaded onto 4-12%. Bis-Tris polyacrylamide gel, followed by electrophoresis in MOPS buffer at 70 V for 10 min, 100 V for 15 min and 150 V for 1h. Transfer was performed with a Biorad Transfer System at 15 V for 30 min. Membranes were blocked in 0.1% TBST/5% milk for 30 min at RT. Immunoblot was carried out with anti-EOLA1 (Novus Biologicals 1:1,000), anti-GAPDH (Santa Cruz, 1:1,000), anti-NAT10 (Santa Cruz 1:1,000), and anti-PABPC1 (Novus, Biologicals 1:1,000) antibodies overnight at 4°C. Membranes were next washed three times in 0.1% TBST for 10 min at RT, followed by a 1h incubation with anti-rabbit antibody (Cell signaling, 1:10,000) or anti-mouse antibody (1:10,000) at RT. After an additional three 10 min washes in 0.1% TBST, GAPDH and NAT10 signals were developed with ECL « Prime » whereas EOLA1 signal was developed with SuperSignal ELISA femto Maximum Sensitivity Substrate (ThermoFisher Scientific). Images were acquired with a ChemiDoc Imaging System (Biorad).

### RNA-seq analysis

Adapters and low-quality bases were removed using Cutadapt (version 1.18). Reads were mapped onto the human reference genome hg38 using STAR and read counts per gene were calculated using RSEM 1.3.1. Differential expressed transcripts were assessed using the R package edgeR.

### Proteomic analysis

Cells were washed twice in 1X cold-PBS and harvested. Cell pellets were lysed and digested using the EasyPep Kit (Thermo A40006). Each sample pellet was resuspended in 200 μL of EasyPep lysis buffer and protein concentration was determined by the BCA method. For each sample 2 μg was treated with 50 μL each of reducing solution and alkylating solution provided with the EasyPep kit, incubated at 25°C for 1hr, then treated with 20 μL of 100 ng/μL Trypsin/LysC (Thermo A40007). Samples were then incubated at 37°C with shaking for 24h after which point 40 μL of 4.2 μg/μL TMTpro 18-plex (Thermo A52045) reagent was added to each sample and incubated for 1h at 25°C with shaking. Excess TMTpro was quenched with 50 μL of 5% hydroxylamine, 20% formic acid for 10 min and samples were then combined. Samples were cleaned using EasyPep Mini columns as described in the manual. Eluted peptides were dried in a speed-vac. LC-MS analysis was performed as in Paul et al., 2025 (30)

### Methanol extraction of nucleosides

#### From cell media

To extract nucleosides from cell media, 50 ul of supernatant was mixed with 450 ul methanol:water solution (8:1 ratio, v:v) at -20°C. The mix was then centrifuged at 4°C for 5min at 16,100g. Supernatants (100uL) were transferred to fresh Eppendorf tubes. Nucleosides were dried for 3h using a rotary vacuum evaporator before being resuspended in 100 uL nuclease-free water.

#### From cytosol

Cells were washed twice with ice cold 1X PBS and harvested in an 800uL methanol:water solution (1:1, v:v). 400 uL of chloroform stored at -20°C was added to the cell suspensions and mixed through rotation at 4°C for 20 min. Phase separation was achieved through centrifugation (5 min-16,100g, -4°C). The upper phase (200 uL) was transferred into a new Eppendorf tube. Nucleosides were dried for 3h using a rotary vacuum evaporator and resuspended in 100uL of nuclease-free water

### Mass-spectrometry analysis of RNA modifications

*In vitro* RNA samples (2 fmol) or RNA isolated from cells (200-500 ng) or free nucleosides were diluted to a total volume of 20 uL with nuclease-free water. A master mix of snake venom phosphodiesterase (0.2 U), calf intestinal phosphatase (2 U) and benzonase (2 U) was prepared with 1 mM MgCl_2_, 5 mM TRIS (pH 8), 10 nmol butylated hydroxytoluene and 5 ug tetrahydrouridine (modified from Heiss et al., 2021(31)) containing isotope analogues of cytidine, N^6^-methyladenosine, 5-methylcytidine and N^4^-acetylcytidine as internal standards. After addition of 15 uL of the mastermix, samples were incubated at 37°C for 2 h. The samples were diluted with 15 uL LC-MS buffer A (0.0075% formic acid in ultrapure water) and filtered for 35 min at 3750g and 4°C through 0.2 um Supor AcroPrep Advance 96-well plates (Pall corporation). Of the filtrate, 39 uL were subjected to mass spectrometric analysis on an Agilent triple quad 6495C mass spectrometer. For HPLC (Agilent 1290 Infinity II), buffer A (described above) and buffer B (0.0075% formic acid in acetonitrile) were used in combination with a alkyl reversed phase column (Agilent Poroshell 120 SB-C8, 2.1 x 150 mm, 2.7 um LC column with the following gradient: 0-2.5 mins 0%, 2.5-3 mins 1%, 3-5 mins 3.5%, 5-7.9 mins 5%, 7.9-8.2 mins 80%, 8.2-11.5 mins 80%, 11.5-12.3 mins 0%, 12.3-14 mins 0%; or Agilent Zorbax RRHD StableBond Aq, 2.1 x 150 mm, 1.8 µm, 80Å with the following gradient: 0-1 mins 0%, 1-1.4 mins 0.1%, 1.4-2.8 mins 0.4%, 2.8-4.2 mins 0.9%, 4.2-5.6 mins 1.6%, 5.6-8 mins 4%, 8-11.5 mins 15%, 15-15.5 mins 50%, 15.5-16.5 mins 50%, 16.5-17 mins 0%, 17-18 mins 0 . All data were analyzed according to published protocol utilizing the isotope dilution technique (32).

### Mass-spectrometry of EOLA1’s interactome and post-translation modifications

Protein LC/MS/MS analysis was carried out using a Thermo Scientific Orbitrap Exploris 240 Mass Spectrometer and a Thermo Dionex UltiMate 3000 RSLCnano System. Peptide mixture from each sample was loaded onto a peptide trap cartridge at a flow rate of 5 μL/min. The trapped peptides were eluted onto a reversed-phase PicoFrit column (New Objective, Woburn, MA) using a linear gradient of acetonitrile (3-36%) in 0.1% formic acid. The elution duration was 110 min at a flow rate of 0.3 μl/min. Eluted peptides from the EasySpray column were ionized and sprayed into the mass spectrometer, using a Nano-EasySpray Ion Source (Thermo) under the following settings: spray voltage, 1.6 kV, Capillary temperature, 275°C. Raw data file acquired from each sample was searched against human protein sequences database using the Proteome Discoverer 1.4 software (Thermo, San Jose, CA) based on the SEQUEST algorithm.

Carbamidomethylation (+57.021 Da) of cysteines was fixed modification, and Oxidation Met and Deamidation Q/N-deamidated (+0.98402 Da), Methyl / +14.016 Da (K, R), HexNAc /+203.079 Da (N, S, T), Acetyl / +42.011 Da (K), Phospho / +79.966 Da (S, T, Y) were set as dynamic modifications. The minimum peptide length was specified to be five amino acids. The precursor mass tolerance was set to 15 ppm, whereas fragment mass tolerance was set to 0.05 Da. The maximum false peptide discovery rate was specified as 0.01. The functions of EOLA1’s partner were visualized through gene ontology (Panther) analysis and STRING-db network analysis.

## Data Availability

High-throughput sequencing datasets generated for this manuscript are publicly available in the Sequence Research Archive (SRA) under accession number SUB15992805. There are no restrictions on any data or materials presented in this paper.

## Results

### EOLA1 and EOLA2 are members of a distinct clade of the ASCH family of PUA-like domains from the bacterial ac^4^C Deacetylase YqfB

Recycling of free ac⁴C through cytidine salvage first requires its deacetylation, as N4-acetylation blocks the initial deamination of cytidine to uridine (16,17). We confirmed this block directly by incubating recombinant human cytidine deaminase (CDA) with ac⁴CTP and analyzing the products by mass spectrometry (Fig. 1a). Whereas unmodified cytidine was efficiently converted to uridine, ac⁴C remained unchanged, with the low level of uridine detected attributable to trace cytidine contamination (Fig. 1a). These results establish that N4-acetylation occludes the exocyclic amine targeted by CDA, implying that free ac⁴C must be deacetylated before it can be salvaged.

**Figure 1.**
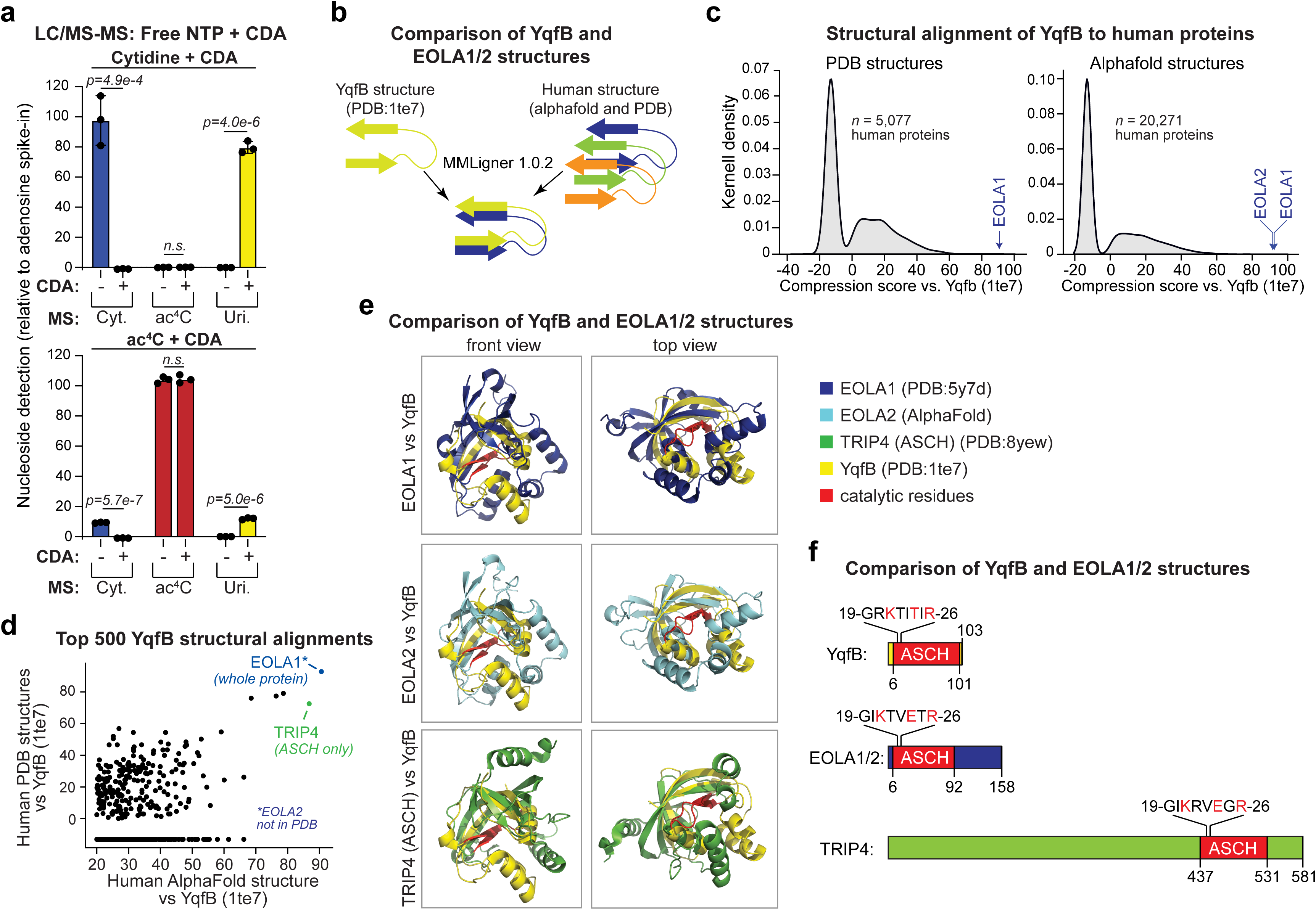
EOLA1 is a human ortholog of YqfB. **(a)** LC-MS/MS analysis of cytidine conversion to uridine upon CDA treatment of ac^4^C and cytidine. Data are mean ± SEM. (n = 3); two-sided unpaired t-test; n.s. denotes not significant (p > 0.05). **(b)** Alignment strategy for comparing the structure of YqfB with structures of ∼20,000 human proteins. **(c)** Distribution of compression scores obtained after aligning the structure of YqfB with structures of human proteins using MMLigner 1.0.2. **(d)** Scatter plot showing the top 500 human proteins with the highest compression scores when aligned to the YqfB structure. **(e)** Structural alignment between YqfB and EOLA1, EOLA2, or TRIP4. **(f)** Comparison of protein domains among YqfB, EOLA1/2, and TRIP4.

Because no human ortholog of the bacterial ac⁴C deacetylase YqfB is detectable by sequence homology, we reasoned that a functional counterpart might instead lie among the multiple, divergent ASCH domain proteins encoded in metazoan genomes (19,24,25) (Table S2). In prokaryotes, contextual associations such as domain architectures and conserved gene-neighborhoods (operons) provide contextual leads regarding the function of poorly characterized proteins(33). For example, in several prokaryotes, genes coding for the YqfB clade of the ASCH domains are linked in a conserved operon with orthologs of TmcA/Nat10 (Fig. S1a), supporting the observation that it operates on ac⁴C generated by this acetyltransferase. We reasoned that comparable contextual associations could help predict ASCH domains that might function as ac⁴C deacetylases in humans. Accordingly, we conducted a systematic sequence search using profile/HMM-based methods to identify ASCH domains in humans and examined the contextual connections within the clade to which they belong. A phylogenetic analysis of the ASCH domains recovered in this search revealed that most eukaryotic ASCH domains belong to a distinct clade from the bacterial YqfB (Fig. S1a). We named this the TRIP4-like clade after the eponymous human protein defining it (19). In addition to TRIP4, this clade also contains two other human proteins, EOLA1 and EOLA2, which, unlike TRIP4, are not multidomain proteins but standalone ASCH domain proteins, just like YqfB (Fig. S1a). Notably, in addition to eukaryotic representatives, the TRIP4-like clade also contains a vast assemblage of bacterial homologs.

Systematic analysis of the prokaryotic versions of the conserved gene-neighborhoods and architectures of the TRIP4-like clade revealed several contextual associations that are indicative of a role in the recognition of modified nucleobases in DNA or RNA (Fig. S1a). In particular, we found several connections to restriction-modification (R-M) systems with modified adenines, as well as systems predicted to generate more complex nucleobase modifications (23,34). ASCH domains encoded by these operons are predicted to either recognize in situ modified bases in nucleic acids or the free nucleotides released upon the degradation of such nucleic acids.

Notably, a further subset of contexts in this clade showed direct fusions or operonic associations of ASCH domains with acetyltransferases of the NAT10/Kre33-like family (Fig. S1a), which are predicted to modify cytosines in DNA or RNA. Hence, we reasoned that members of the TRIP4- like clade could also potentially recognize or operate on ac⁴C, as has been reported for members of the YqfB clade. This, in turn, led us to focus on EOLA1 and EOLA2 as they are standalone versions of the ASCH domain comparable to YqfB (18–22).

Pairwise structural alignments between the NMR solution structure of YqfB (PDB: 1TE7) (35) and AlphaFold-predicted human protein structures using MMLigner (28) produced negative compression scores for most alignments, indicating no significant similarity (Figs. 1b, 1c).

However, it yielded EOLA1 and EOLA2 as the highest-scoring significant matches (Figs. 1c, 1d). We next aligned the YqfB NMR structure to the experimentally solved EOLA1 X-ray crystal structure (PDB: 5Y7D) (22). This revealed complete collinearity between YqfB and EOLA1 in the region encompassing the proposed catalytic residues in the former: K21 and R26 that are essential for YqfB deacetylase activity are conserved as is in EOLA1/2 (Figs. 1e, 1f).

The third predicted active site residue, two positions upstream of R26, is a glutamate (E24) in EOLA1/EOLA2, which is absolutely conserved across the Trip4-like clade. In contrast, in the YqfB clade, it is an absolutely conserved threonine/serine. Nevertheless, the polar side chains are similarly positioned in both clades, suggesting they could participate similarly in the deacetylation reaction (Fig. 1f).

#### EOLA1/2 deletion results in increased free ac^4^C levels

We next asked whether EOLA1 and EOLA2 function in ac⁴C metabolism in human cells. Both genes are ubiquitously expressed at low levels across tissues, and both transcripts are associated with translating polysomes in HeLa cells (Figs. S2a, S2b). The two loci share extensive nucleotide similarity, consistent with a recent duplication event (Fig. S2c) (36). This similarity and genomic proximity preclude the selective deletion of one paralog. Accordingly, we used a CRISPR/Cas9 strategy targeting a conserved coding region in EOLA1 to simultaneously disrupt both loci and yield a double knockout cell line (Fig. 2a). Loss of EOLA1/2 was confirmed at the DNA, RNA, and protein levels by PCR, RT–qPCR, and Western blotting using reagents that recognize both paralogs (Figs. 2b-2d). Disruption of both genes was further validated by Nanopore sequencing (Fig. S2d). In contrast, the intervening genes *TMEM185A* and *MAGEA11* remained intact by genomic DNA PCR (Fig. 2c), indicating that downstream phenotypes arise from the loss of EOLA1/2 rather than a disruption of the local genomic context.

**Figure 2.**
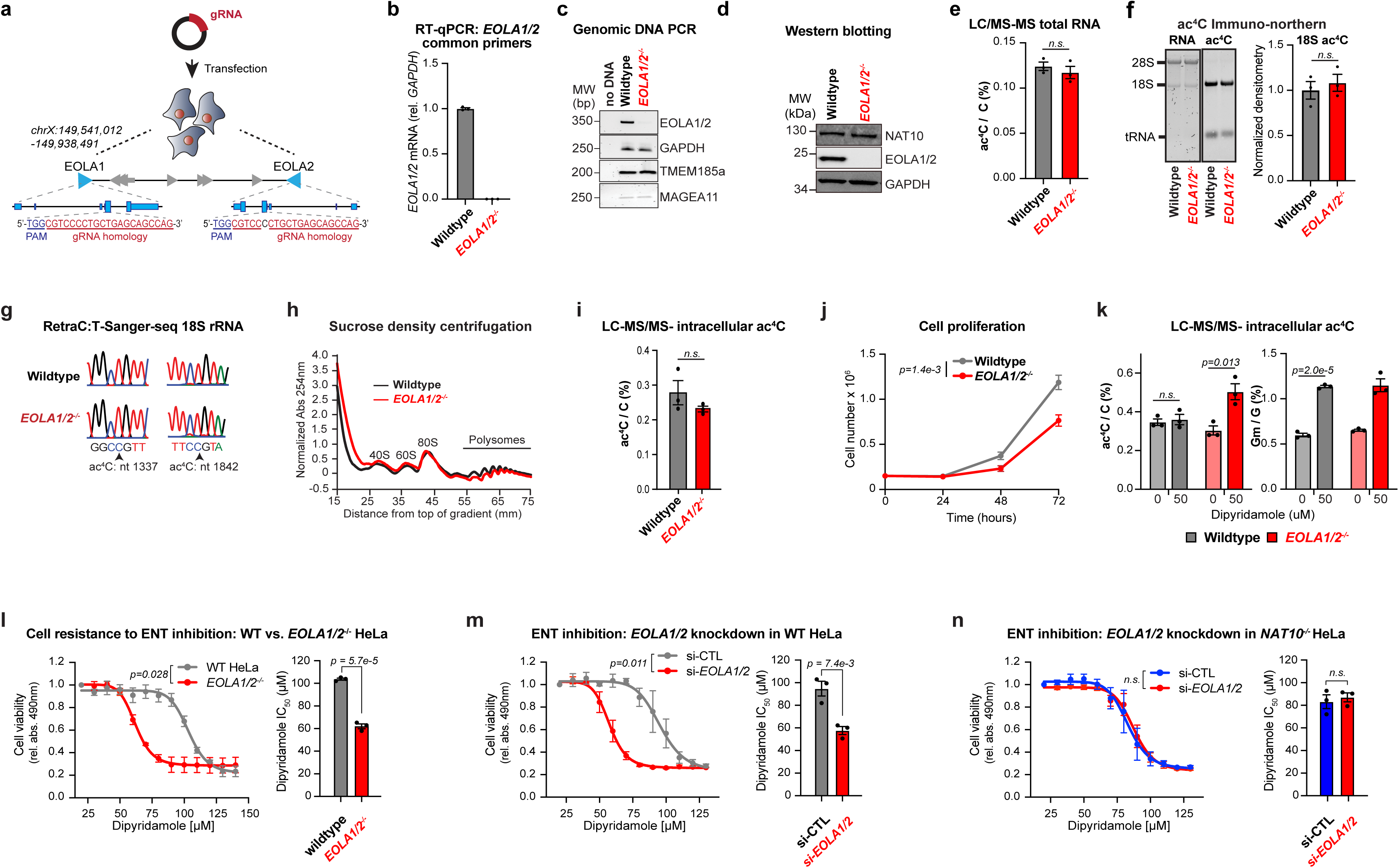
EOLA1/2 deletion results in increased free ac^4^C levels. **(a)** Schematic of the CRISPR-Cas9 strategy to delete EOLA1/2. **(b)** RT-qPCR quantification of *EOLA1/2* mRNA in EOLA1/2 knockout HeLa. Data are mean ± SEM (n = 3). **(c)** Genomic PCR validating deletion specificity at the *EOLA1/2* locus, with amplification of *GAPDH* and the intervening genes *MAGEA11* and *TMEM185A* shown as controls. **(d)** Validation of protein ablation in *EOLA1/2^-/-^* HeLa by Western blot. **(e)** LC–MS/MS quantification of ac⁴C levels in total RNA from wildtype and *EOLA1/2^-/-^* HeLa. Data represent mean ± SEM (n = 3); two-sided unpaired t-test; n.s. denotes p > 0.05. **(f)** Immuno-northern blot detection of ac⁴C in total RNA from wildtype and *EOLA1/2^-/-^* HeLa. Representative blots from three independent experiments are shown (left), with densitometric quantification of ac⁴C levels in 18S rRNA (right). Data represent mean ± SEM (n = 3); two-sided unpaired t-test; n.s. denotes p > 0.05. **(g)** RetraC:T-seq quantification of 18S rRNA acetylation at positions 1337 and 1842 in wildtype and *EOLA1/2^-/-^* HeLa. Sanger traces are representative of three independent experiments. **(h)** Sucrose gradient profile in wildtype and *EOLA1/2^-/-^* HeLa. Profile is representative of three independent experiments. **(i)** LC-MS/MS quantification of free ac^4^C extracted from the cytosol of wildtype and *EOLA1/2^-/-^* HeLa. Data are mean ± SEM (n = 3); n.s. denotes p > 0.05; two-sided unpaired t-test. **(j)** Comparison of cell proliferation in wildtype and *EOLA1/2^-/-^* HeLa. Data are mean ± SEM (n = 3); two-way ANOVA. **(k)** LC-MS/MS quantification of intracellular free ac^4^C in wildtype and *EOLA1/2^-/-^* HeLa treated with dipyridamole (DPM) to inhibit nucleoside export. G_m_ serves as a positive control for inhibition efficiency. Data are mean ± SEM (n = 3); n.s. denotes p > 0.05; two-sided unpaired t-test. **(l–n)** HeLa cell viability in response to increasing DPM in wild-type versus EOLA1/2 KO cells (l), or following control versus EOLA1/2 silencing in wild-type (m) and NAT10 KO (n) cells. Dose–response curves represent four-parameter logistic (variable-slope) nonlinear regression fits, with significance assessed by two-way repeated-measures ANOVA; corresponding IC₅₀ values are shown as bar plots and compared by two-sided unpaired Student’s t-test. Data are mean ± SEM (n = 3); n.s., not significant (p > 0.05).

Consistent with the inability of YqfB to deacetylate ac⁴C within intact RNA (37), EOLA1/2 loss did not alter ac⁴C abundance in established RNA pools. LC–MS/MS analysis of digested total RNA revealed no difference in ac^4^C levels between wild-type and EOLA1/2-null cells, while the control modifications m⁶A and m⁵C were likewise unchanged (Figs. 2e, S2e). Analysis of purified small and poly(A) RNA similarly showed no effect on ac⁴C in tRNA- and mRNA-enriched fractions, respectively (Figs. S2f, S2g). These findings were corroborated by immuno- northern blotting with an ac⁴C-specific antibody and by RetraC:T, which detects ac⁴C as C-to-T mismatches following chemical reduction and reverse transcription (29), at the two established ac⁴C sites in 18S rRNA (Figs. 2f, 2g). In keeping with preserved rRNA and tRNA function, polysome profiles were indistinguishable between genotypes (Fig. 2h). Thus, as predicted from the substrate specificity of YqfB, EOLA1/2 deletion does not measurably perturb ac⁴C in canonical RNA substrates.

Because free ribonucleotides released during RNA turnover are subsequently salvaged or exported, we asked whether EOLA1/2 regulate ac⁴C in the soluble nucleotide pool, analogous to the in vitro activity of YqfB toward free ac⁴C (18). Free nucleotides were extracted from intracellular and extracellular fractions, with enrichment for non-polymerized species confirmed by the absence of detectable rRNA (Fig. S3a). At steady state, intracellular ac⁴C, m⁵C, and 2′-O- methylguanosine (Gm) were indistinguishable between wild-type and *EOLA1/2⁻/⁻* cells (Figs. 2i, S3b). Despite this apparent lack of a molecular phenotype, *EOLA1/2⁻/⁻* HeLa cells proliferated more slowly than wild-type cells (Fig. 2j). This suggested that EOLA1/2 loss produces a defect not readily captured under steady-state conditions. Notably, ac⁴C was readily detected in the extracellular fraction, whereas the abundant RNA modifications m⁶A and m⁵C were indistinguishable from background (Fig. S3c), suggesting that preferential export may mask intracellular ac⁴C accumulation.

To test this, we blocked equilibrative nucleoside transport with dipyridamole (DPM), thereby retaining free nucleosides within the cell (2). Effective transport inhibition was confirmed by increased intracellular Gm in both genotypes (Fig. 2k). Under these conditions, free ac⁴C accumulated specifically in *EOLA1/2⁻/⁻* cells while remaining unchanged in wild-type cells (Fig. 2k). This accumulation was accompanied by reduced viability of *EOLA1/2⁻/⁻*cells relative to wild-type cells following DPM treatment (Figs. 2l). Comparable DPM sensitivity was observed following siRNA-mediated co-depletion of EOLA1 and EOLA2 in wild-type cells, with knockdown confirmed by RT–qPCR and Western blotting (Figs. 2m, S3d, S3e), indicating that the phenotype was not specific to the knockout genotype. By contrast, responses to sodium arsenite and hydrogen peroxide were comparable across genotypes (Figs. S3f, S3g), demonstrating that EOLA1/2 loss does not confer a general sensitivity to cellular stress.

Notably, because ac⁴C is present at very low levels relative to unmodified cytidine, impaired ac⁴C recycling would not be expected to appreciably deplete the cytidine pool. Indeed, cytidine levels were unchanged in EOLA1/2-null cells (Fig. S3h). This raises the possibility that the associated fitness defects arise from aberrant accumulation of free ac⁴C rather than from reduced cytidine availability for salvage. Supporting this interpretation, EOLA1/2 depletion did not alter DPM sensitivity in NAT10-null cells, which lack ac⁴C synthesis (Fig. 2n). The absence of the phenotype in NAT10-null cells firmly links the viability defect to ac⁴C metabolism.

Although the mechanism connecting altered ac⁴C turnover to cell fitness remains to be established, further characterization revealed mild basal phospho-γH2AX accumulation in EOLA1/2-null cells, consistent with low-level genotoxic stress (Fig. S3i). RNA-seq and quantitative proteomics also identified changes in pathways related to mitochondrial respiration, ATP production, purine catabolism, and other aspects of nucleotide metabolism (Figs. S3j-m; Tables S3, S4). Together with the proliferation delay, these findings indicate that chronic EOLA1/2 loss is accompanied by broader changes in cellular metabolism and nucleotide homeostasis, whereas export blockade reveals a specific ac⁴C-dependent vulnerability.

#### EOLA1 overexpression limits free ac⁴C accumulation

Having established that EOLA1/2 loss promotes free ac⁴C accumulation when nucleoside export is blocked, we asked whether restoring EOLA1 expression would reverse this phenotype. To this end, EOLA1 and EOLA2 were expressed as C-terminal FLAG fusions to preserve the N-terminal catalytic region and enable protein detection and recovery (Table S1). Although RT– qPCR confirmed increased expression of both transcripts, only EOLA1-FLAG was readily detected by Western blotting (Fig. S4a), suggesting that EOLA2 may be comparatively unstable. The two paralogs differ at only two residues, W53C and V112A, which map to adjacent α-helices in structural models and may contribute to their differential expression (Fig. S4b). We therefore focused subsequent analyses on EOLA1.

To specifically interrogate EOLA1 enzymatic activity, we additionally generated a catalytically inactive mutant (R26A), based on prior studies showing that the equivalent residue is essential for YqfB deacetylase function (18) (Fig. 3a). Wildtype and R26A EOLA1-FLAG constructs were transiently expressed in HeLa cells and verified by Western blotting (Fig. 3b). Importantly, EOLA1 overexpression did not alter the expression of NAT10, confirming that any changes in acetylation were not attributable to changed levels of the ac⁴C writer (Fig. 3b).

**Figure 3.**
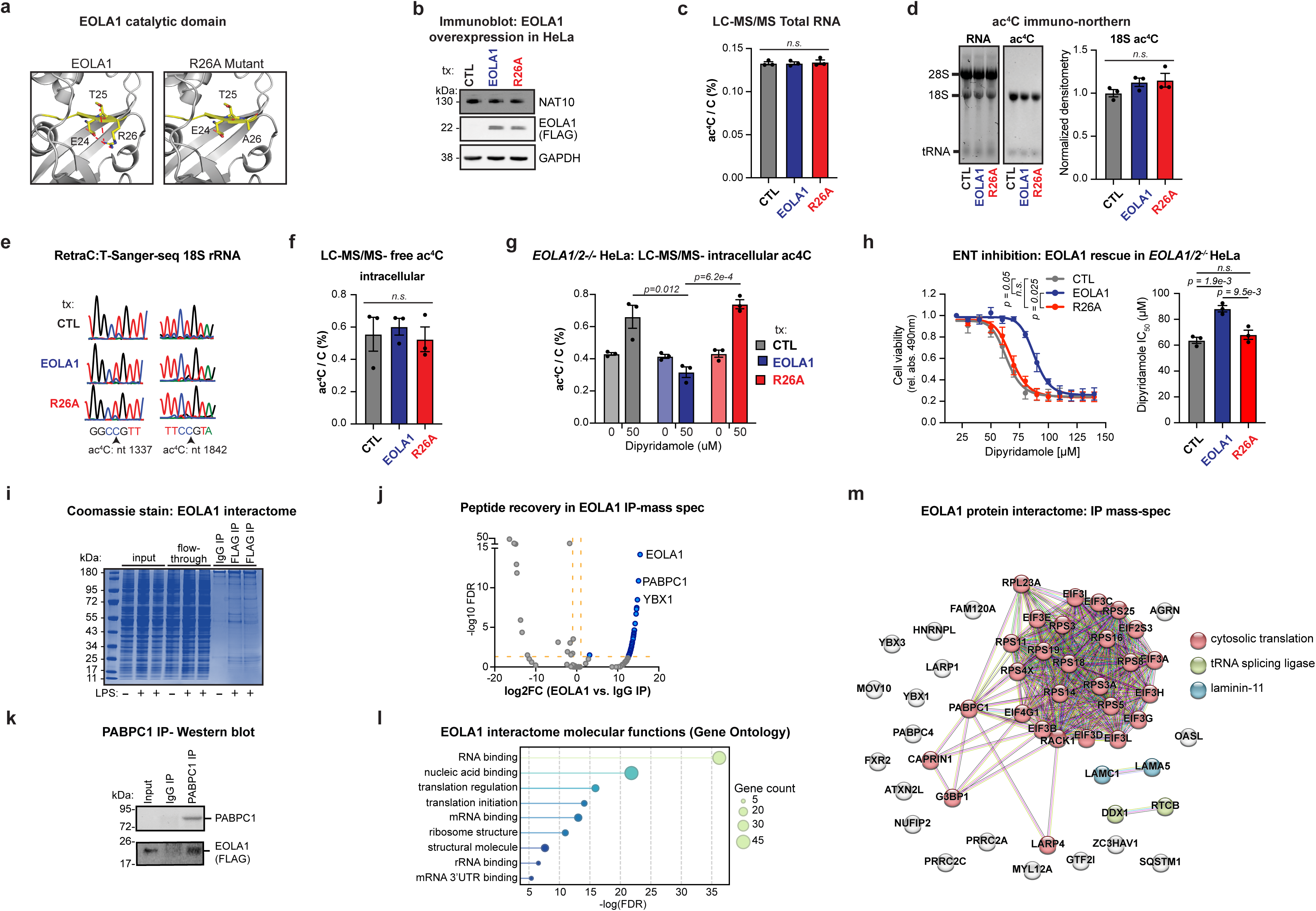
EOLA1 overexpression results in decreased free ac^4^C levels. **(a)** Structural representation of the catalytic site of wild-type EOLA1 and the R26A catalytic mutant. **(b)** Western blot validation of wildtype and R26A mutant EOLA1 overexpression. Images are representative of three biological replicates. **(c)** LC-MS/MS quantification of RNA acetylation in EOLA1 rescued *EOLA1/2^-/-^* HeLa. Data are mean ± SEM (n = 3); two-sided unpaired t-test; n.s. denotes p > 0.05. **(d)** Immuno-northern blot quantification of total RNA acetylation in EOLA1 rescued *EOLA1/2^-/-^* HeLa. Images are representative of three independent experiments. Bar plots show mean ± SEM (n = 3); two-sided unpaired t-test; n.s. denotes p > 0.05. **(e)** RetraC:T-seq quantification of 18S rRNA acetylation upon EOLA1 overexpression in *EOLA1/2^-/-^* HeLa. Sequencing traces are representative of three independent experiments. **(f)** LC-MS/MS quantification of intracellular free ac^4^C upon EOLA1 overexpression in *EOLA1/2^-/-^* HeLa. Data are mean ± SEM (n = 3); two-sided unpaired t-test; n.s. denotes p > 0.05. **(g)** LC-MS/MS quantification of intracellular free ac^4^C in DPM-treated *EOLA1/2^-/-^* HeLa with EOLA1 rescue. Data are mean ± SEM (n = 3); two-sided unpaired t-test, only significant values (p > 0.05) are shown. **(h)** Cell viability in response to increasing DPM in EOLA1-rescued *EOLA1/2^-/-^* HeLa. Dose– response curves represent four-parameter logistic (variable-slope) nonlinear regression fits, with significance assessed by two-way repeated-measures ANOVA with Holm-corrected pairwise comparisons; corresponding IC₅₀ values are shown as bar plots and compared by two-sided unpaired Student’s t-test. Data are mean ± SEM (n = 3); n.s., not significant (p > 0.05). **(i)** Coomassie staining of EOLA1-interacting proteins following FLAG immunoprecipitation of EOLA1. **(j)** Quantitative mass spectrometry analysis of EOLA1-interacting proteins. **(k)** Validation of EOLA1-PABPC1 interaction by co-immunoprecipitation and Western blot. **(l)** Molecular function Gene Ontology analysis of EOLA1-interacting proteins. **(m)** Network analysis of EOLA1-interacting proteins.

Consistent with observations in EOLA1/2-deleted cells, EOLA1 overexpression did not alter ac⁴C levels within canonical RNA substrates. MS analysis showed that neither wildtype nor R26A EOLA1-FLAG changed ac4C levels in total or small RNA, nor did they affect other RNA modifications such as m⁶A or m⁵C (Figs. 3c, S4c–d). Likewise, ac⁴C immuno-northern blotting and RetraC:T-seq revealed no differences in acetylation at the established 18S rRNA or tRNA sites (Figs. 3d, 3e, S4e, S4f), confirming that EOLA1 does not influence ac⁴C installation in cellular RNA. MS analysis of purified free nucleotides further demonstrated that EOLA1 overexpression did not alter intracellular ac⁴C or the control modification m⁵C, nor extracellular ac⁴C, at steady state (Figs. 3f, S4g–i).

In contrast, exogenous EOLA1 restored intracellular ac⁴C homeostasis when nucleoside export was inhibited in *EOLA1/2^-/-^* HeLa cells. Following DPM treatment, cells expressing vector or the catalytically inactive R26A mutant accumulated intracellular free ac⁴C, whereas reconstitution with wildtype EOLA1 fully suppressed this accumulation (Fig. 3g). Intracellular free Gm increased similarly in wildtype- and R26A-expressing cells, confirming comparable inhibition of nucleoside transport (Fig. S4j). Strikingly, the same catalytic requirement was observed at the cellular level: wildtype EOLA1 restored DPM resistance, whereas R26A failed to rescue viability (Fig. 3h). The concordant rescue of ac⁴C accumulation and cell fitness by wildtype, but not catalytically inactive, EOLA1 establishes that its deacetylase activity is required to maintain ac⁴C homeostasis when nucleoside export is restricted. These findings further demonstrate that the phenotypes associated with EOLA1/2 loss reflect the absence of catalytic activity rather than secondary adaptation of the knockout cells.

Expression of tagged EOLA1 also enabled examination of its protein interactome by affinity purification and MS. Because EOLA1 has previously been linked to endothelial responses to lipopolysaccharide (LPS) (38), interactomes were compared in untreated and LPS-treated cells. FLAG-EOLA1 was immunoprecipitated under non-denaturing conditions, and Coomassie-stained SDS–PAGE revealed discrete bands enriched relative to input and IgG controls (Fig. 3i). No significant differences were detected between untreated and LPS-treated conditions using thresholds of FDR < 0.05 and log₂FC > 1 (Fig. S4k), indicating that LPS did not measurably remodel the EOLA1 interactome under these conditions. By contrast, comparison with IgG controls identified 50 proteins significantly enriched with EOLA1 at FDR < 0.05 (Fig. 3j; Table S5). Reciprocal immunoprecipitation of the top interactor, PABPC1, followed by EOLA1 immunoblotting confirmed the association and validated the affinity-purification approach (Fig. 3k).

Markedly, Gene Ontology analysis of the co-purified proteins revealed strong enrichment for nucleotide-metabolism pathways, with nearly all interactors annotated as RNA-binding proteins (Fig. 3l). High-confidence STRING-db network analysis further showed that more than half of the interactors participate in mRNA translation and identified additional enrichment for tRNA-processing factors (Fig. 3m). Collectively, these findings place EOLA1 within ribonucleoprotein machineries associated with the two principal cellular reservoirs of ac⁴C, 18S rRNA and tRNA, positioning it in proximity to the RNA pools from which free ac⁴C is likely released during turnover.

#### EOLA1 deacetylates free and structurally accessible ac⁴C

Characterization of EOLA1 activity in HeLa cells suggested a role in the deacetylation of free ac⁴C. To directly establish EOLA1 catalytic activity, we purified FLAG-tagged wildtype and R26A mutant EOLA1 from HeLa cells under mildly denaturing conditions to minimize co-purification of associated proteins. Recovery and purity were verified by SDS-PAGE, Western blotting, and Coomassie staining (Figs. 4a, S5a). Incubation of purified EOLA1 with free ac⁴CTP revealed robust deacetylase activity: LC-MS/MS with isotopically labeled standards showed a clear reduction in ac⁴C and a compensatory increase in cytidine in the presence of wildtype, but not mutant, EOLA1 (Fig. 4b). Levels of unmodified CTP, m⁶ATP, and m⁵CTP were unaffected by EOLA1, establishing its specificity for ac⁴C (Figs. 4b, S5b).

**Figure 4.**
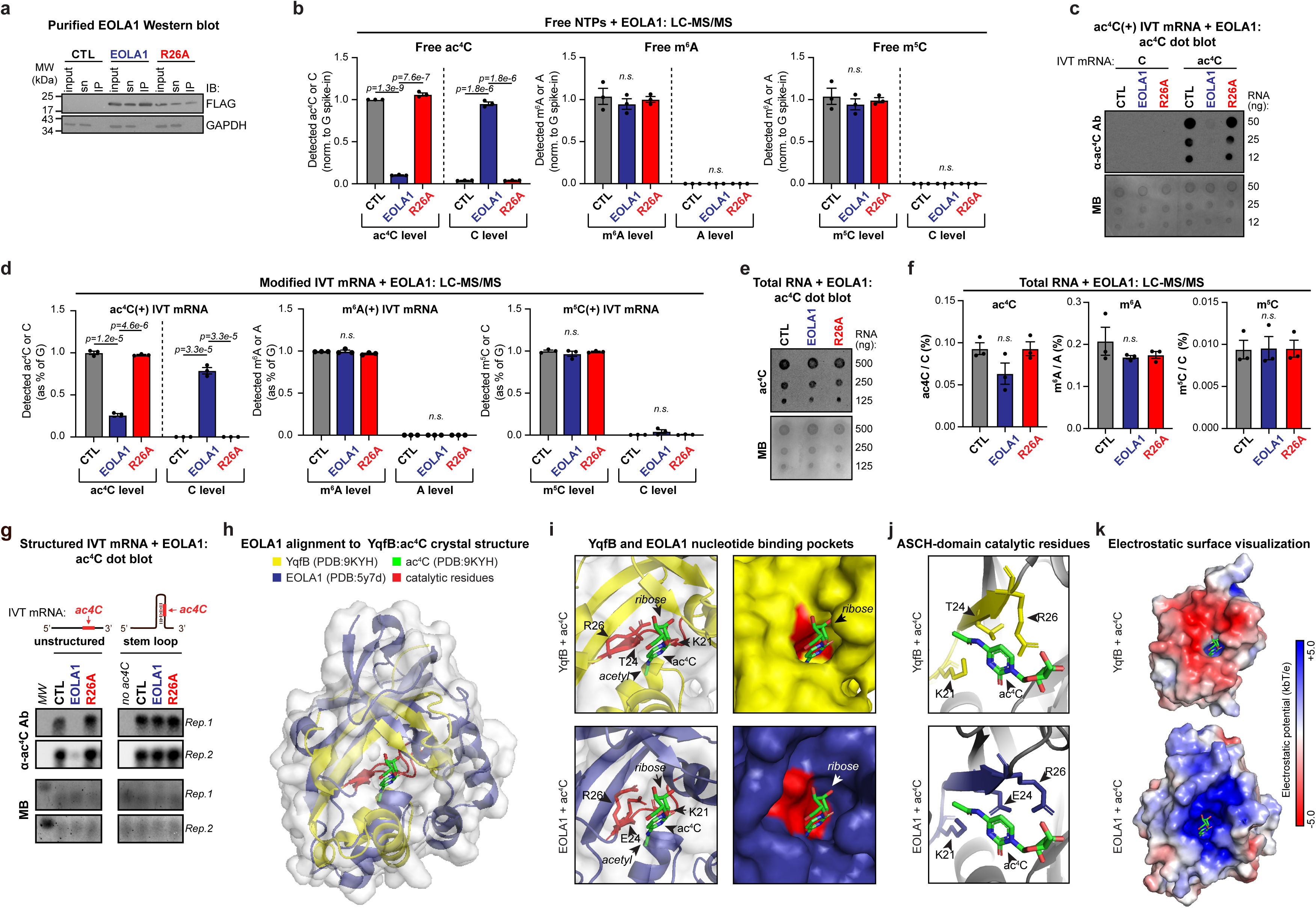
EOLA1 deacetylates ac^4^C in vitro. **(a)** Western blot of purified wildtype and R26A mutant EOLA1. **(b)** LC-MS/MS quantification of free ac^4^C, m^6^A, and m^5^C after in vitro incubation with wildtype or R26A mutant EOLA1. Data are mean ± SEM (n = 3); two-sided unpaired t-test; n.s. denotes p > 0.05 **(c)** Dot blot for ac^4^C in acetylated *FLuc* IVT RNA after incubation with wildtype or R26A mutant EOLA1. Images are representative of three independent experiments. **(d)** LC-MS/MS quantification of ac^4^C, m^6^A, and m^5^C in *FLuc* IVT RNA incorporating the modifications as indicated, after incubation with purified wildtype or mutant EOLA1. Data are mean ± SEM (n = 3); two-sided unpaired t-test; n.s. denotes p > 0.05. **(e)** Dot blot for ac^4^C in HeLa total RNA after incubation with wildtype or mutant EOLA in vitro. Images are representative of three independent experiments. **(f)** LC-MS/MS quantification of ac4C levels in HeLa total RNA after incubation with wildtype or mutant EOLA1 in vitro. Data are mean ± SEM (n = 3); two-sided unpaired t-test; n.s. denotes p > 0.05. **(g)** Immuno-northern blot for ac^4^C in matched unstructured and hairpin-structured acetylated IVT RNA after incubation with wildtype or R26A mutant EOLA1. Individual replicates are shown. **(h)** Structural alignment of EOLA1 with YqfB and ac^4^C docking. ac^4^C is shown in green; catalytic residues are shown in red. **(i)** Magnified view of the nucleotide-binding pocket from the EOLA1 structural alignment with YqfB docked to ac^4^C. **(j)** Magnified view of the catalytic residues from the EOLA1 structural alignment with YqfB docked to ac^4^C. **(k)** Electrostatic surface visualization of the nucleotide-binding pocket of YqfB and EOLA1 in the presence of ac^4^C.

Together with the cellular data showing altered free ac⁴C levels following modulation of EOLA1 (Figs. 2 and 3), these findings establish EOLA1 as a human deacetylase that acts on free ac⁴C. To determine whether EOLA1 could also act on ac⁴C incorporated into RNA in this minimal in vitro system, we generated an in vitro-transcribed (IVT) mRNA reporter encoding Firefly Luciferase (FLuc) using either unmodified NTPs or ac⁴CTP in place of CTP. Given the pronounced association of EOLA1 with PABPC1 in the proteomic analysis, the IVT mRNAs were tested with or without capping and polyadenylation (Fig. S5c). Surprisingly, in both contexts, dot blot analysis revealed a loss of ac⁴C signal following incubation with wildtype, but not mutant, EOLA1 (Figs. 4c, S5d). Consistently, LC-MS/MS confirmed a marked reduction in ac⁴C and reciprocal cytidine accumulation only in the presence of wildtype protein (Fig. 4d). By contrast, unmodified IVT mRNA and transcripts incorporating m⁶A or m⁵C were unaffected by EOLA1 (Figs. 4d, S5e, S5f).

EOLA1-catalyzed deacetylation of ac⁴C within synthetic mRNAs contrasted with our cellular findings, in which neither loss nor overexpression of EOLA1 altered ac⁴C levels in its endogenous RNA substrates (Figs. 2e, 3c). This difference was maintained in vitro: ac⁴C within total RNA isolated from HeLa cells remained refractory to EOLA1-catalyzed deacetylation by both dot blot and LC-MS/MS analysis (Figs. 4e, 4f). Thus, incorporation into RNA does not inherently prevent EOLA1 activity; rather, ac⁴C within its endogenous substrates is selectively protected from deacetylation, raising the question of the molecular basis for this distinction.

We reasoned that RNA secondary structure might contribute to this differential susceptibility. Cellular ac⁴C is concentrated at defined positions within highly structured rRNAs and tRNAs, where the modified base participates in stable helical interactions. By contrast, uniform substitution of CTP with ac⁴CTP in an IVT transcript generates numerous modified sites distributed across a range of structural environments, including accessible regions. To test whether local structure governs EOLA1 activity, we generated matched IVT RNA substrates in which ac⁴C-containing sequences were designed to adopt either structured or unstructured conformations. The structured substrate incorporated ac⁴C into a hairpin used previously with the intended conformations confirmed by mFold (11,39). Purified wildtype EOLA1, but not the R26A mutant, efficiently deacetylated ac⁴C in the unstructured substrate, whereas the modification was largely protected when embedded within the structured RNA (Fig. 4g). These findings identify local RNA structure as one determinant limiting EOLA1 activity toward RNA-incorporated ac⁴C. Notably, the bacterial homolog YqfB efficiently deacetylates free ac⁴C in vitro but lacks intrinsic RNA-binding activity (18,37), suggesting that limited engagement of RNA substrates may be a broader feature of ASCH-domain YqfB homologs.

To explore the mechanistic basis limiting EOLA1 activity against cellular RNA, we examined the solved crystal structure of YqfB bound to ac⁴C (PDB: 9KYH) (40). In this assembly, the acetyl group of ac⁴C occupies a narrow pocket of YqfB encompassing the catalytic triad (R26, T24, K21), while the ribose is oriented outward and partly exposed to the solvent (Figs. 4h-j). Alignment of EOLA1 with this structure revealed a similar architecture: the acetyl group extends into a confined pocket where the cytidine ring and side chain form contacts with catalytic residues R26, E24, and K21 (Figs. 4h-j). This positioning is consistent with the acetyl carbonyl group engaging the positively charged environment of the catalytic pocket through dipolar interactions that stabilize substrate orientation for hydrolysis. The structural profile thus supports the premise that the physiological substrate of EOLA1 is free ac⁴C. The activity observed against in vitro–transcribed RNA likely reflects enhanced accessibility of synthetic molecules, which are relatively flexible and devoid of bound proteins, enabling base flipping into the catalytic pocket. In contrast, ac⁴C within rRNA and tRNA is engaged in Watson–Crick base-pairing within helical stems, where the nucleobase is buried in the duplex and inaccessible to the catalytic pocket, providing a rationale for the limited activity of EOLA1 on cellular RNA. Both YqfB and EOLA1 display nucleotide-binding pockets characterized by positive electrostatic potential, consistent with their interaction with the negatively charged phosphate backbone of RNA. In EOLA1, this positive surface extends beyond the narrow catalytic cleft (Fig. 4k), suggesting an electrostatically guided docking mechanism that facilitates transient interactions with RNA during substrate search or engagement.

Notably, a recent structural study comparing ASCH-domain proteins did not detect deacetylation of free ac⁴C by murine EOLA1 (mEOLA1) in vitro (40). To address this discrepancy, we compared the corresponding crystal structures and found that the two proteins are highly conserved, providing no obvious structural basis for their differing activities (Fig. S5h).

However, mEOLA1 was produced in *E. coli*, whereas the human EOLA1 used in our assays was purified from transfected HeLa cells. Because bacterially expressed proteins often lack key post-translational modifications (PTMs), we reasoned that PTM status could influence catalytic function. Mass spectrometric analysis of human EOLA1 purified from HeLa cells revealed abundant phosphorylation at serine 7 (S7), accounting for approximately 44% of peptides spanning this residue (Table S6). This position is universally occupied by an alcoholic residue, serine or threonine, within the Trip4-like clade and is also conserved in a subset of the YqfB clade, including YqfB itself. Located within the catalytic pocket and oriented toward three core active-site residues (Figs. S5h–i), this hydroxyl-bearing residue may contribute to substrate recognition or catalysis. Thus, the apparent discrepancy between human and murine EOLA1 activity may reflect differences in post-translational modification rather than intrinsic biochemical divergence. These findings establish human EOLA1 as a bona fide ac⁴C deacetylase and raise the possibility that its catalytic efficiency is regulated by phosphorylation of a conserved active-site residue.

#### EOLA1 does not deacetylate ac^4^C within therapeutic mRNAs

We recently reported that ac⁴C represents a viable addition to the toolbox of modifications used in mRNA-based medicines, limiting immune stimulation comparably to the industry-standard N1-methylpseudouridine (m¹Ψ) while enhancing translation fidelity (15). The ability of EOLA1 to deacetylate IVT mRNA in vitro raised the possibility that it might also act on synthetic mRNAs in vivo, potentially blunting their translational benefit. To test this, LNP-encapsulated ac⁴C-, m¹Ψ-, and unmodified NanoLuc mRNAs were delivered into wildtype and *EOLA1/2^-/-^* HeLa cells (Figs. 5a, S6a). Transfection efficiencies were comparable by RT-qPCR, and translational output was quantified by luminescence (Figs. 5b, 5c). As observed previously (15), ac⁴C-modified mRNA supported the highest NanoLuc protein output (Fig. 5c). Importantly, loss of EOLA1/2 did not selectively increase luminescence from ac⁴C-modified mRNA. Instead, all three transcripts showed a similar modest increase in output in *EOLA1/2^-/-^*cells, indicating a small, general effect of genotype on translation rather than one specific to ac⁴C (Fig. 5c).

**Figure 5.**
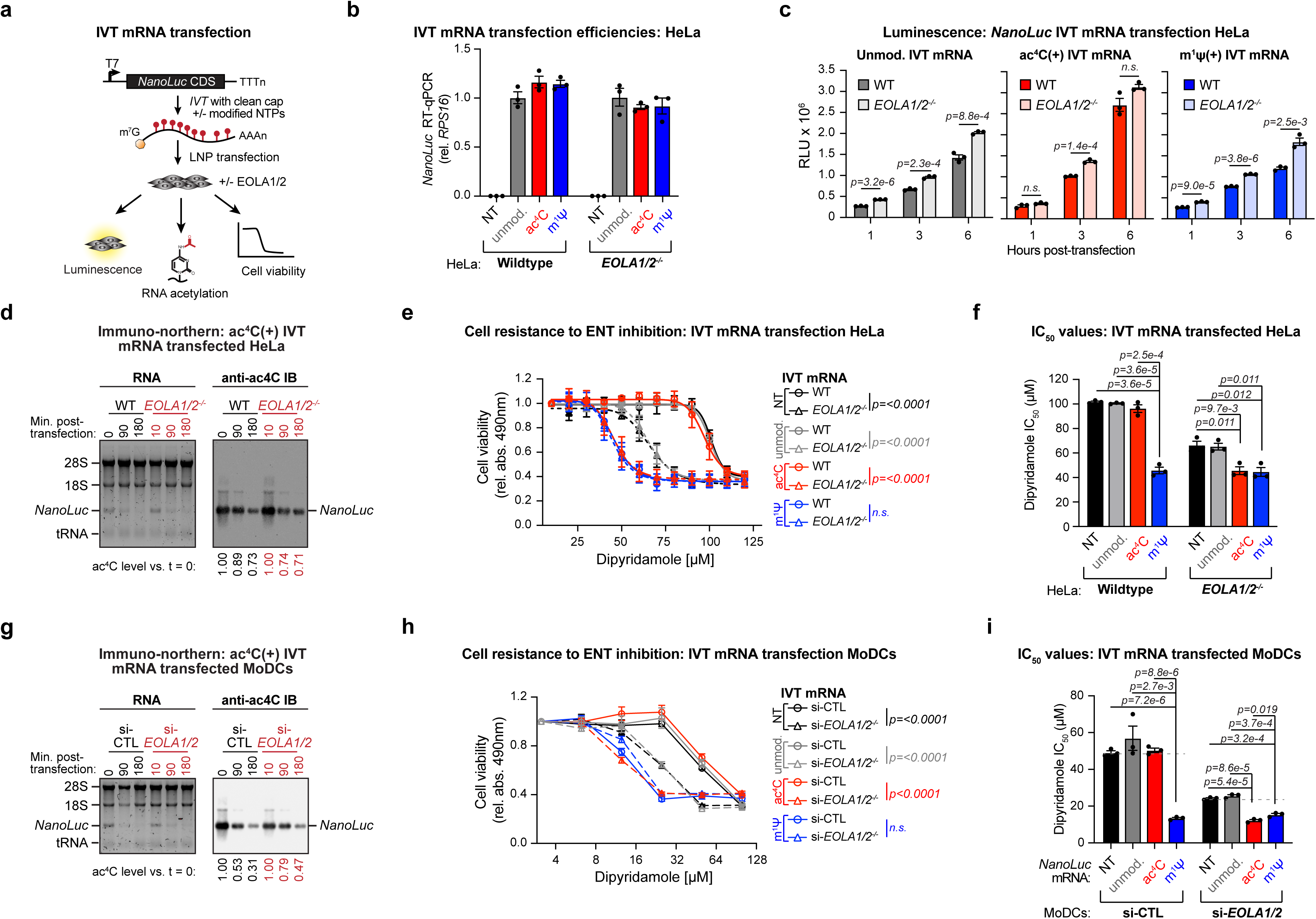
EOLA1 does not deacetylate ac^4^C within therapeutic mRNAs. **(a)** Schematic of IVT mRNA production and transfection. **(b)** RT-qPCR assessment of *NanoLuc* transfection efficiencies in wildtype and *EOLA1/2^-/-^* HeLa. Data are mean ± SEM (n = 3). **(c)** Luminescence from wildtype and *EOLA1/2^-/-^* HeLa transfected with ac^4^C-modified, m^1^Ψ-modified or unmodified *NanoLuc* mRNA. Data are mean ± SEM (n = 3); two-sided unpaired t-test; n.s. denotes p > 0.05. **(d)** ac^4^C immuno-northern blot of total RNA from wildtype and *EOLA1/2^-/-^*HeLa transfected with ac^4^C-modified *NanoLuc* mRNA. *NanoLuc* acetylation levels determined by densitometry are shown below the image. **(e)** Cell resistance to DPM in wildtype and *EOLA1/2^-/-^* HeLa transfected with modified or unmodified *NanoLuc* mRNA. Data are mean ± SEM (n = 3); two-way ANOVA. **(f)** DPM IC₅₀ in wildtype and *EOLA1/2^-/-^* HeLa transfected with modified or unmodified*NanoLuc* mRNA. Data are mean ± SEM (n = 3); two-sided unpaired t-test. **(g)** ac^4^C immuno-northern blot of total RNA from control and EOLA1/2 depleted MoDCs transfected with ac^4^C-modified *NanoLuc* mRNA. *NanoLuc* acetylation levels determined by densitometry. **(h)** Cell resistance to DPM in control and EOLA1/2 depleted MoDCs transfected with modified or unmodified *NanoLuc* mRNA. Data are mean ± SEM (n = 3); two-way ANOVA. **(i)** DPM IC₅₀ in control and EOLA1/2 depleted MoDCs transfected with modified or unmodified *NanoLuc* mRNA. Data are mean ± SEM (n = 3); two-sided unpaired t-test.

The absence of an ac⁴C-specific difference between wildtype and *EOLA1/2^-/-^* HeLa cells suggests that the deacetylase does not target the mRNA polymer in vivo. This distinction from the minimal in vitro system may reflect the more complex cellular environment, in which RNA structure and RNA-binding protein networks limit EOLA1/2 access. Consistent with this interpretation, immuno-northern analysis of RNA isolated from transfected cells showed that ac⁴C levels in *NanoLuc* mRNA were unchanged between wildtype and EOLA1/2-null HeLa cells (Fig. 5d).

Because IVT mRNA transfection substantially increases the intracellular pool of modified nucleotides, we next asked whether these nucleotides alter sensitivity to ENT blockade. Wildtype and EOLA1/2-null HeLa cells were transfected with ac⁴C-, m¹Ψ-, or unmodified *NanoLuc* mRNAs and exposed to a range of DPM concentrations. Cell viability assays revealed several patterns. First, m¹Ψ-modified mRNA markedly increased sensitivity to DPM relative to ac⁴C- modified or unmodified mRNA (Figs. 5e, 5f). Although the basis of this increased toxicity is unknown, m¹Ψ is not a natural modification in human RNA and may therefore lack dedicated salvage pathways, leaving export as its primary route of clearance. Second, in wildtype HeLa cells, ac⁴C-modified and unmodified mRNAs behaved similarly to non-transfected controls, indicating that canonical nucleotides and ac⁴C are effectively metabolized when EOLA1/2 activity is present (Figs. 5e, 5f). By contrast, EOLA1/2-null cells exhibited elevated DPM sensitivity across conditions, consistent with accumulation of ac⁴C from endogenous sources. However, whereas unmodified mRNA increased DPM sensitivity to the same extent as in non-transfected knockout controls, ac⁴C-modified mRNA caused an additional decrease in viability that matched the m¹Ψ profile (Fig. 5e). These results support a model in which EOLA1/2 deacetylate free ac⁴C released during IVT mRNA turnover, thereby preventing its accumulation and mitigating toxicity during ENT blockade.

These findings were recapitulated in primary human monocyte-derived dendritic cells (MoDCs), the principal cell type responsible for antigen uptake in mRNA-based vaccine platforms (41).

Monocytes were isolated from peripheral blood, differentiated into MoDCs, and treated with either control siRNA (si-CTL) or siRNA targeting a shared region of EOLA1 and EOLA2 (si- *EOLA1/2*) (Figs. S6b–d). As in HeLa cells, immuno-northern analysis showed that ac⁴C levels in transfected *NanoLuc* mRNA were similar between si-CTL and si-*EOLA1/2* MoDCs (Fig. 5g), confirming that EOLA1/2 do not deacetylate the mRNA polymer in this primary cell type.

Viability measurements during DPM treatment also mirrored the trends observed in HeLa cells. m¹Ψ-modified mRNA substantially increased sensitivity to ENT inhibition, whereas ac⁴C- modified and unmodified mRNAs behaved similarly to non-transfected controls in si-CTL MoDCs (Figs. 5h, 5i). In EOLA1/2-deficient MoDCs, DPM sensitivity was elevated and was most pronounced following transfection with ac⁴C-modified mRNA, reaching levels observed with m¹Ψ-modified mRNA (Figs. 5h, 5i). Together, these results show that EOLA1/2 do not substantially impact ac⁴C levels within IVT mRNAs, but instead act on free ac⁴C generated during mRNA turnover. Overall, the discovery of a dedicated ac⁴C recycling mechanism, together with the metabolic liabilities observed for m¹Ψ, supports the use of ac⁴C in next-generation mRNA therapeutics.

## Discussion

RNA therapeutics introduce substantial quantities of chemically modified nucleosides into cells, yet comparatively little is known about how these building blocks are handled after the RNA is degraded. Our findings identify EOLA1 as an ac⁴C deacetylase that enables cytidine released from ac⁴C-modified RNA to re-enter pyrimidine salvage. This activity may represent an intrinsic metabolic advantage of ac⁴C in therapeutic mRNAs: the modification remains stable during the functional lifetime of the RNA but can be removed after RNA turnover, allowing the underlying cytidine to be recycled. More broadly, our results show that the regulation of RNA modifications extends beyond RNA molecules themselves and can also occur at the level of free modified nucleosides.

We identify EOLA1, and likely its paralog EOLA2, in carrying out this processing step. Because N4-acetylation blocks the deamination of cytidine to uridine that initiates salvage (16), free ac⁴C cannot be recycled directly and must first be exported or deacetylated. EOLA1/2 activity therefore serves two related functions: it returns cytidine to the salvage pool as an energetically favorable alternative to de novo synthesis and limits the intracellular accumulation of an otherwise poorly metabolized modified nucleoside. Under basal conditions, ac⁴C appears to be cleared efficiently, but conditions that increase its intracellular abundance, including the elevated RNA metabolism associated with some cancers, may increase reliance on this pathway (42,43).

Structural alignment of EOLA1 with ac⁴C-bound YqfB provides a basis for this substrate selectivity. The acetylated base inserts into a narrow catalytic pocket formed by K21, E24, and R26 and recessed from the nucleotide-binding surface (44,45). Productive catalysis would therefore require the ac⁴C base to rotate out of the RNA and into the pocket (Fig. 4h). Base flipping is a recurrent strategy among nucleic-acid-modifying enzymes, including DNA methyltransferases, glycosylases, and pseudouridine synthases (46–49), and the PUA-like ASCH fold occurs in several domains that engage target bases in a flipped-out conformation (23,50).

Because extrusion from a stable helix is energetically costly, free ac⁴C is more readily accommodated than ac⁴C embedded within structured RNA (51). This model is consistent with our observation that EOLA1 deacetylates free ac⁴C but not ac⁴C paired within the helical stems of rRNA and tRNA (Fig. 4g). Beyond the pocket, an extended electropositive surface surrounding the active site (Fig. 4k) may contribute to substrate engagement while restricting productive catalysis to accessible bases.

The ASCH domain shared by EOLA1/2 and YqfB defines a conserved framework linking RNA metabolism to the processing of modified nucleoside byproducts (18,19,21,22). Across eukaryotes, members of this clade are fused to RNA-associated modules, including the PWI and Zn-cluster RNA-binding domains of TRIP4, an AlkB-related 2-oxoglutarate- and Fe-dependent dioxygenase, an S1-like RNA-binding domain, and a HAD-superfamily phosphatase predicted to act on RNA termini and degradation products (Fig. S1a) (52,53). These architectures support the view that ASCH-domain proteins act on or near RNA. Consistent with this, EOLA1 co-purifies with numerous RNA-associated proteins, suggesting that protein interactions may position it near sites of RNA turnover where free ac⁴C is generated. In bacteria, close homologs are frequently encoded in restriction-modification operons alongside endoRNases predicted to cleave tRNAs (34,54) (Fig. S1a). Given the prevalence of ac⁴C in bacterial tRNAs, these ASCH proteins may process modified nucleosides released during tRNA degradation and could thereby contribute to phage defense.

EOLA1 activity may also be regulated. We find prominent phosphorylation at S7 near the N-terminus (Table S6), a conserved residue oriented toward the catalytic pocket, where modification could influence catalysis or substrate engagement (Table S6). EOLA1 abundance is also regulated under inflammatory conditions, as its expression increases following LPS exposure role (38). These findings point to coordinated regulation of EOLA1 activity and expression under conditions that alter ac⁴C burden, including bacterial infection, where degradation of ac⁴C-containing bacterial tRNAs introduces this modified nucleoside into host cells (8,55).

In summary, we identify EOLA1 as a human ac⁴C deacetylase that acts on the free nucleoside rather than on ac⁴C embedded within intact RNA, and implicate EOLA2 in the same pathway.

This activity returns ac⁴C-derived cytidine to pyrimidine salvage after RNA turnover, linking RNA degradation to modified-nucleoside homeostasis. This distinction may be particularly advantageous for therapeutic mRNAs: ac⁴C remains stable within the functional transcript but can be processed after RNA turnover, potentially limiting metabolic burden without eroding its translational benefits.

## Acknowledgments

We thank members of the Center for Cancer Research Genomics Core for providing Illumina sequencing services. We thank members of CCR Protein Characterization Laboratory (PCL) for providing global comparative proteomic services. This work used the computational resources of the National Institutes of Health (NIH) HPC Biowulf cluster (http://hpc.nih.gov). This work is supported by the Intramural Research Program of NIH, the Center for Cancer Research, the National Cancer Institute and the National Library of Medicine. The contributions of the NIH author(s) are considered Works of the United States Government. The findings and conclusions presented in this paper are those of the authors and do not necessarily reflect the views of the NIH or the U.S. Department of Health and Human Services.

## Author contributions

S.R., and S.O.: conceptualization; S.R., A.M.B., L.A., and S.O.: methodology; S.R., S.S., M.P., N.T., C.A., A.R., and H.B.: data analysis and curation; S.R.: investigation and validation; S.R.: writing original draft; S.R., and S.O.: writing review and editing; S.O.: supervision; S.O.: funding acquisition.

---

**Figure S1.**
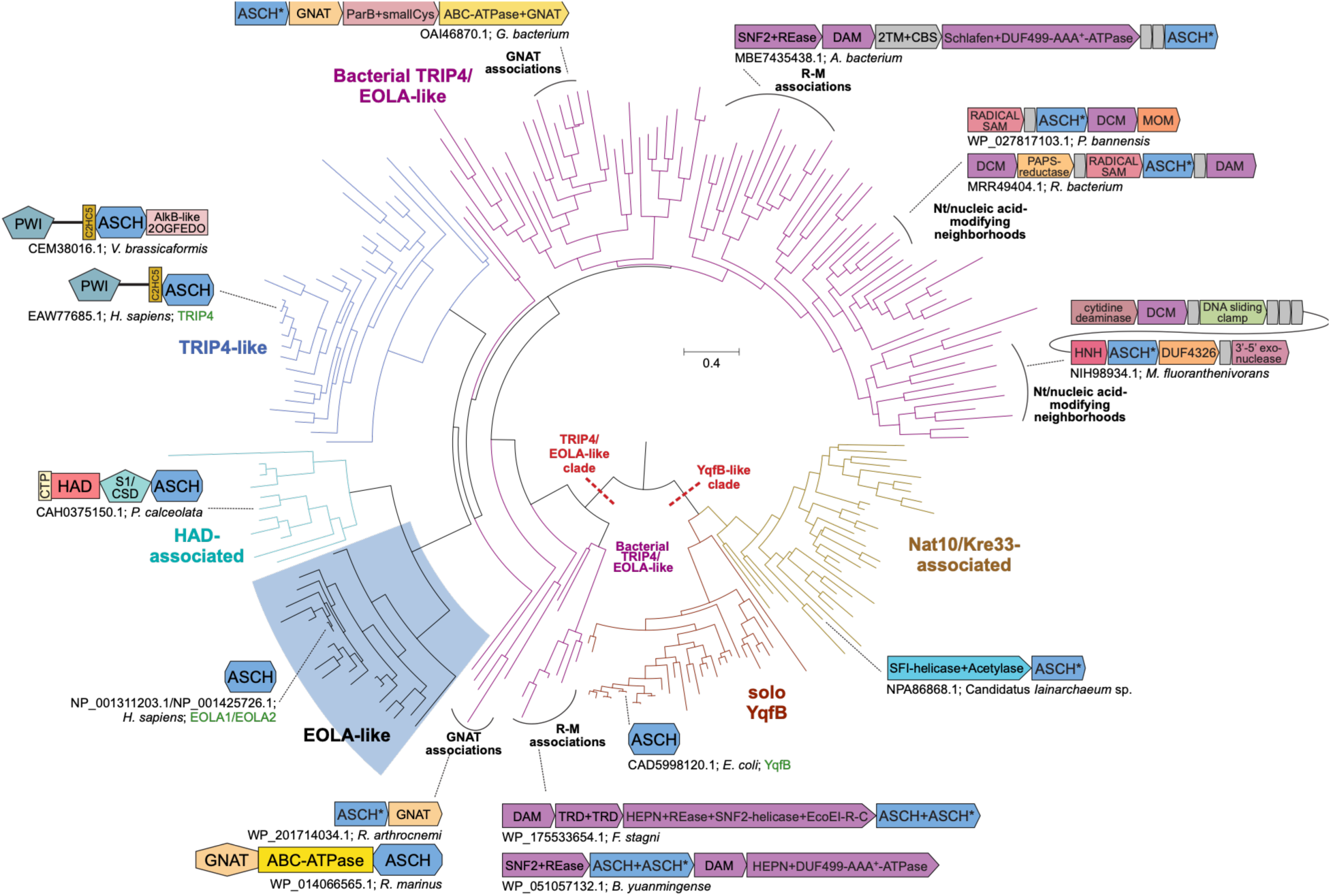
Phylogenetic tree depicting relationships between clades in the ASCH superfamily. **(a)** Branches are colored according to their predominant architectural associations, with the background of the EOLA-like lineage shaded blue. The two major ASCH monophyletic clades are labeled and marked by dashed red lines. Genome contexts of representative sequences are depicted as box arrows in the case of gene neighborhoods and adjoined shapes in the case of individual protein domain architectures. In these contexts, the ASCH domain and the genes encoding it are colored in blue. Selected sequences are connected to their relevant branch(es) by dashed lines.

**Figure S2.**
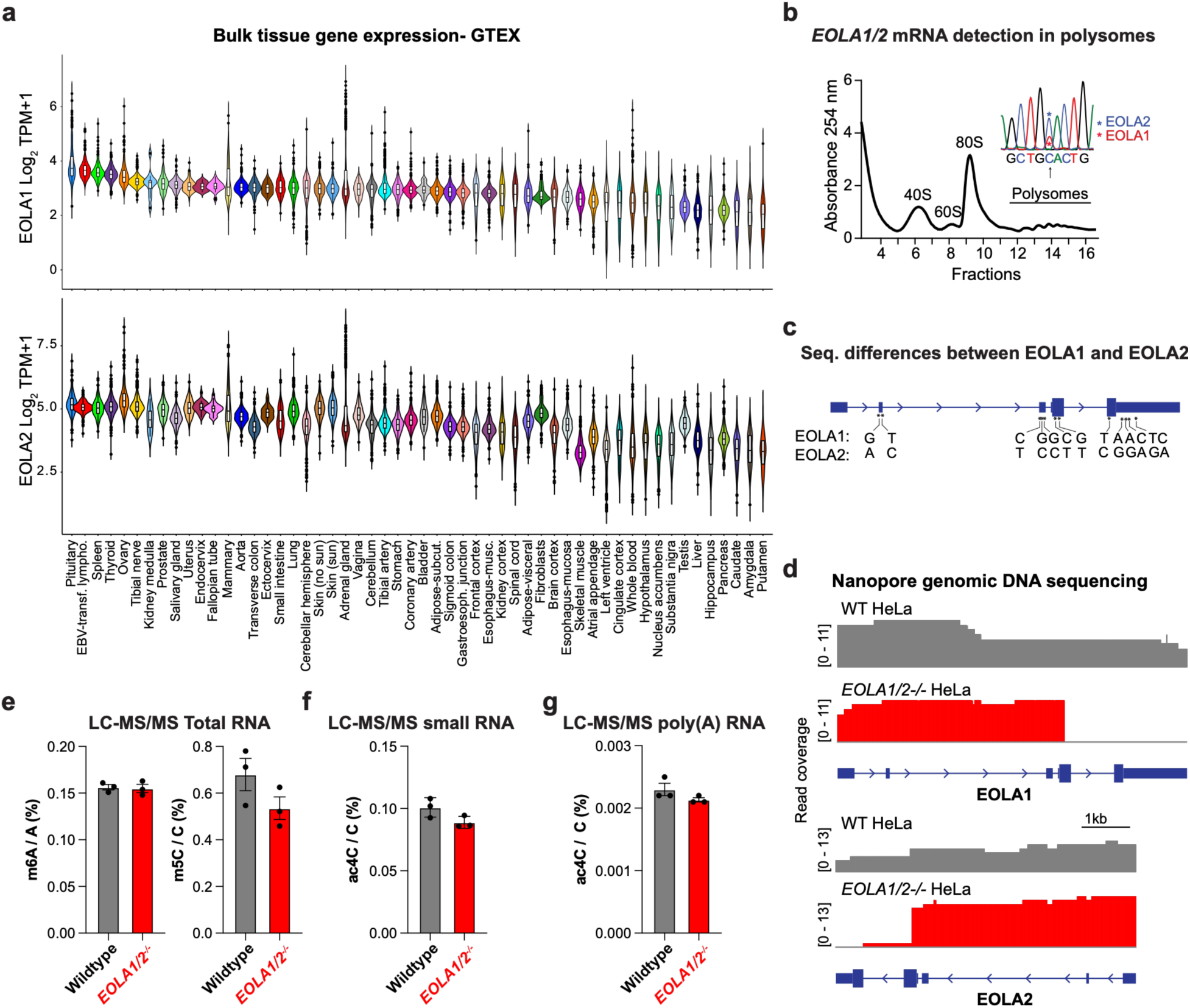
EOLA1/2 deletion in HeLa. **(a)** GTEx mRNA expression of EOLA1 and EOLA2 across human tissues. **(b)** Detection of EOLA1 and EOLA2 mRNAs in polysome-associated fractions by Sanger sequencing. **(c)** Nucleotide sequence differences between EOLA1 and EOLA2. **(d)** Read-depth drop at CRISPR–Cas9 cut sites detected by nanopore sequencing of genomic DNA. **(e-g)** LC–MS/MS of m^6^A, m^5^C or ac^4^C in total, small or poly(A) RNA from wildtype and *EOLA1/2^-/-^* HeLa, as indicated. Data are mean ± SEM (n = 3); two-sided unpaired t-test; not significant (n.s.) throughout.

**Figure S3.**
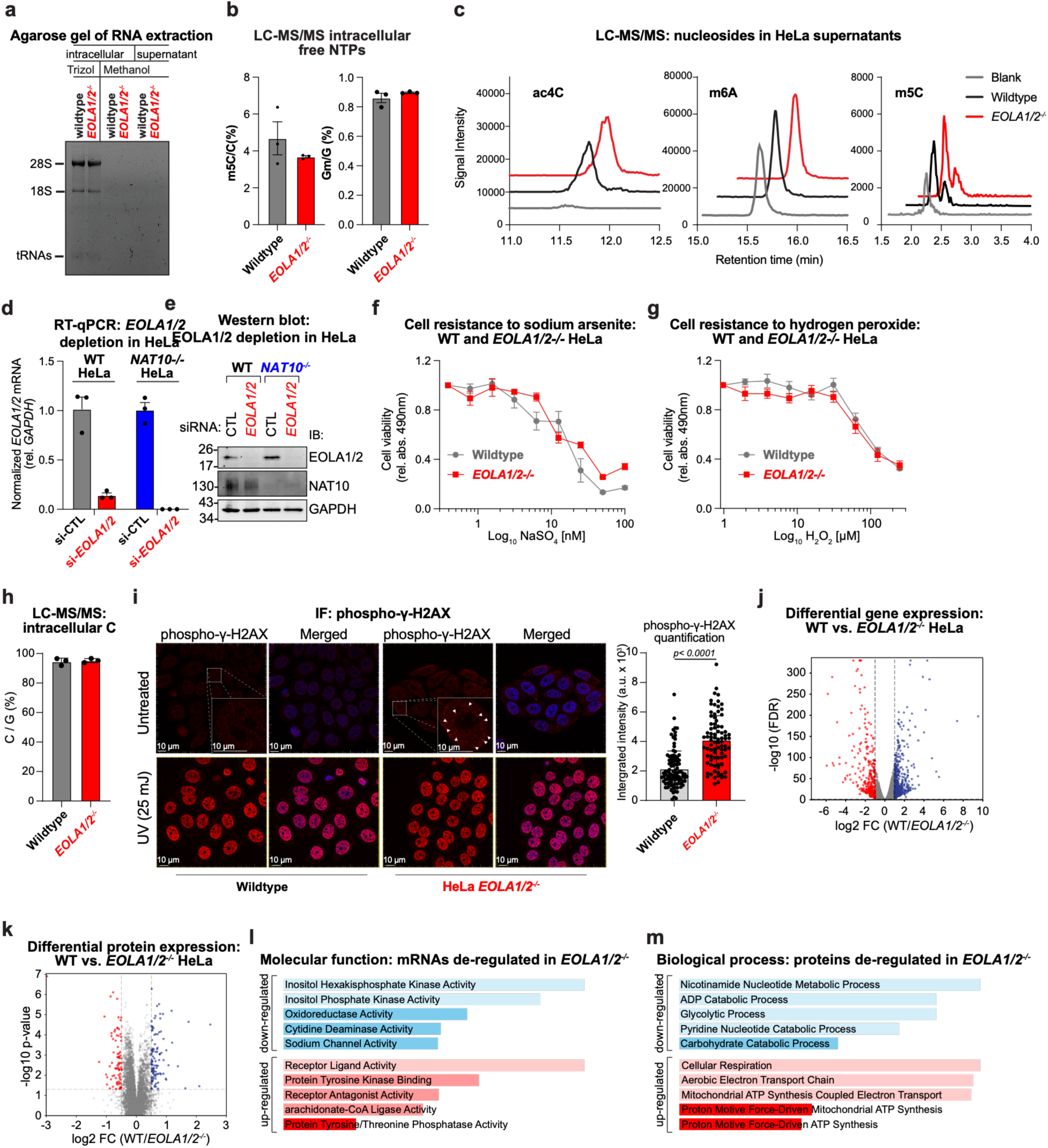
Characterization of EOLA1/2 deleted HeLa. **(a)** RNA contamination assessment of methanol-extracted nucleosides by RNA electrophoresis; TRIzol-extracted RNA shown as positive control. **(b)** LC–MS/MS for intracellular m^5^C and Gm in wildtype and *EOLA1/2^-/-^* cells. **(c)** Representative LC–MS/MS chromatograms of free ac^4^C, m6A, and m5C detected in culture media from wild-type and EOLA1/2 knockout cells. (**d)** RT–qPCR validation of EOLA1/2 silencing in HeLa wildtype and *NAT10^-/-^* HeLa. Data are mean ± SEM (n = 3). **(e)** Immunoblot analysis of EOLA1/2 following silencing in HeLa wildtype and *NAT10^-/-^* cells. **(f,g)** Cell viability of *EOLA1/2^-/-^* cells following treatment with sodium arsenite (f) or hydrogen peroxide (g). Data are mean ± SEM (n = 3). Overall resistance calculated through area under the curve (AUC) across all concentrations (log-scale) for each replicate; unpaired Student’s t-test, n.s. **(h)** LC-MS/MS for intracellular free cytidine in wildtype and *EOLA1/2^-/-^* HeLa. Data are mean ± SD (n=3) **(i)** Representative images and quantification of phospho-ɣH2AX staining in wildtype and *EOLA1/2^-/-^* cells. UV treated cells are positive controls for phospho-ɣH2AX staining. Horizontal white bars represent 10μm scales. Bar plot represents mean ± SD of 83 WT nuclei and 99 *EOLA1/2^-/-^* nuclei; unpaired T-test with Welch correction. **(j)** Volcano plot of RNA-seq analysis comparing mRNA expression in wildtype and *EOLA1/2^-/-^*HeLa. Red dots represent downregulated mRNAs (log₂FC < -1, FDR < 0.05); blue dots represent upregulated mRNAs (log₂FC > 1, FDR < 0.05); n = 3 biological replicates per condition. **(k)** Volcano plot of proteome expression analysis in wildtype and *EOLA1/2^-/-^* HeLa. Red dots represent downregulated proteins (log₂FC < -0.5, p-value < 0.05); blue dots represent upregulated proteins (log₂FC > 0.5, p-value < 0.05); n = 3 biological replicates per condition. *EOLA1/2^-/-^*cells compared to wildtype. **(l)** Molecular function GO term analysis of differentially expressed mRNAs in *EOLA1/2^-/-^* HeLa. Under- and over-represented gene sets are shown in blue and red, respectively. **(m)** Biological process GO term analysis of dysregulated proteins in *EOLA1/2^-/-^*HeLa. Under- and over-represented gene sets are shown in blue and red, respectively.

**Figure S4.**
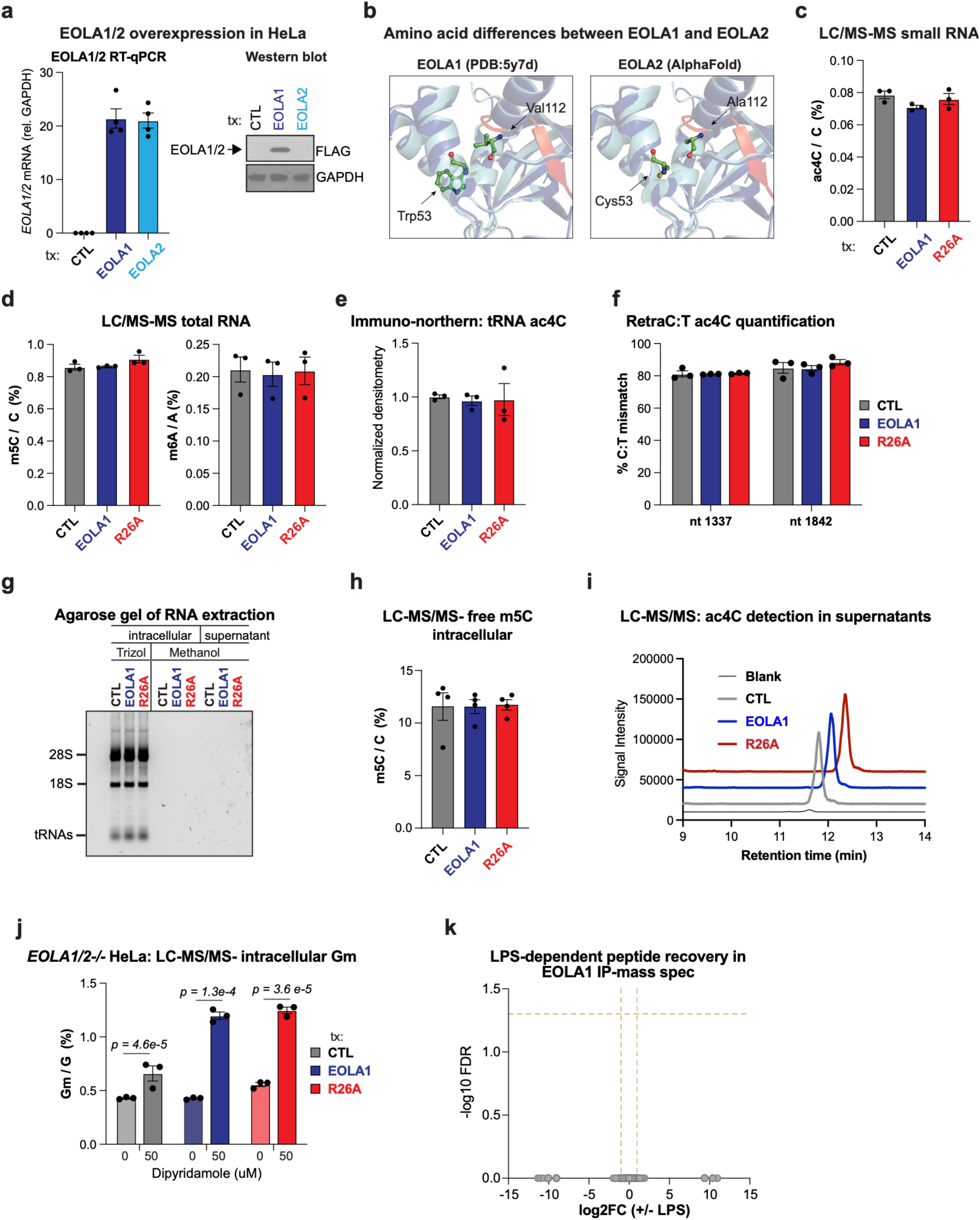
EOLA1 overexpression in HeLa. **(a)** RT–qPCR and immunoblot validation of EOLA1 and EOLA2 overexpression. **(b)** Structural comparison of EOLA1 and EOLA2 at mismatch positions. **(c-d)** LC–MS/MS quantification of ac4C in small RNA following overexpression of EOLA1 or the R26A catalytic mutant. LC–MS/MS of ac^4^C, m^6^A, or m^5^C in small or total RNA following overexpression of EOLA1 or the R26A catalytic mutant, as indicated. Data are mean ± SEM (n = 3); two-sided unpaired t-test; n.s. throughout. **(e)** Immuno-northern analysis of tRNA acetylation following EOLA1 overexpression. Data are mean ± SEM (n = 3); two-sided unpaired t-test; n.s. **(f)** RetraC:T-seq quantification of ac^4^C sites upon EOLA1 overexpression. Data are mean ± SEM (n = 3); two-sided unpaired t-test; n.s. **(g)** RNA contamination assessment of methanol-extracted nucleosides by RNA electrophoresis; TRIzol-extracted RNA shown as positive control. **(h)** LC–MS/MS quantification of intracellular free m^5^C following EOLA1 overexpression. Data are mean ± SEM (n = 3); two-sided unpaired t-test; n.s. **(i)** Representative LC–MS/MS chromatograms of free ac^4^C detected in culture media upon EOLA1 overexpression. **(j)** LC–MS/MS quantification of intracellular free 2′-O-methylguanosine (Gm) following EOLA1 overexpression with or without dipyridamole treatment. Data are mean ± SEM (n = 3); two-sided unpaired t-test; only significant values are shown. **(k)** LPS-specific EOLA1 protein interactors identified by mass spectrometry using CPM-normalized spectral counts and Fisher’s exact test with Benjamini–Hochberg correction.

**Figure S5.**
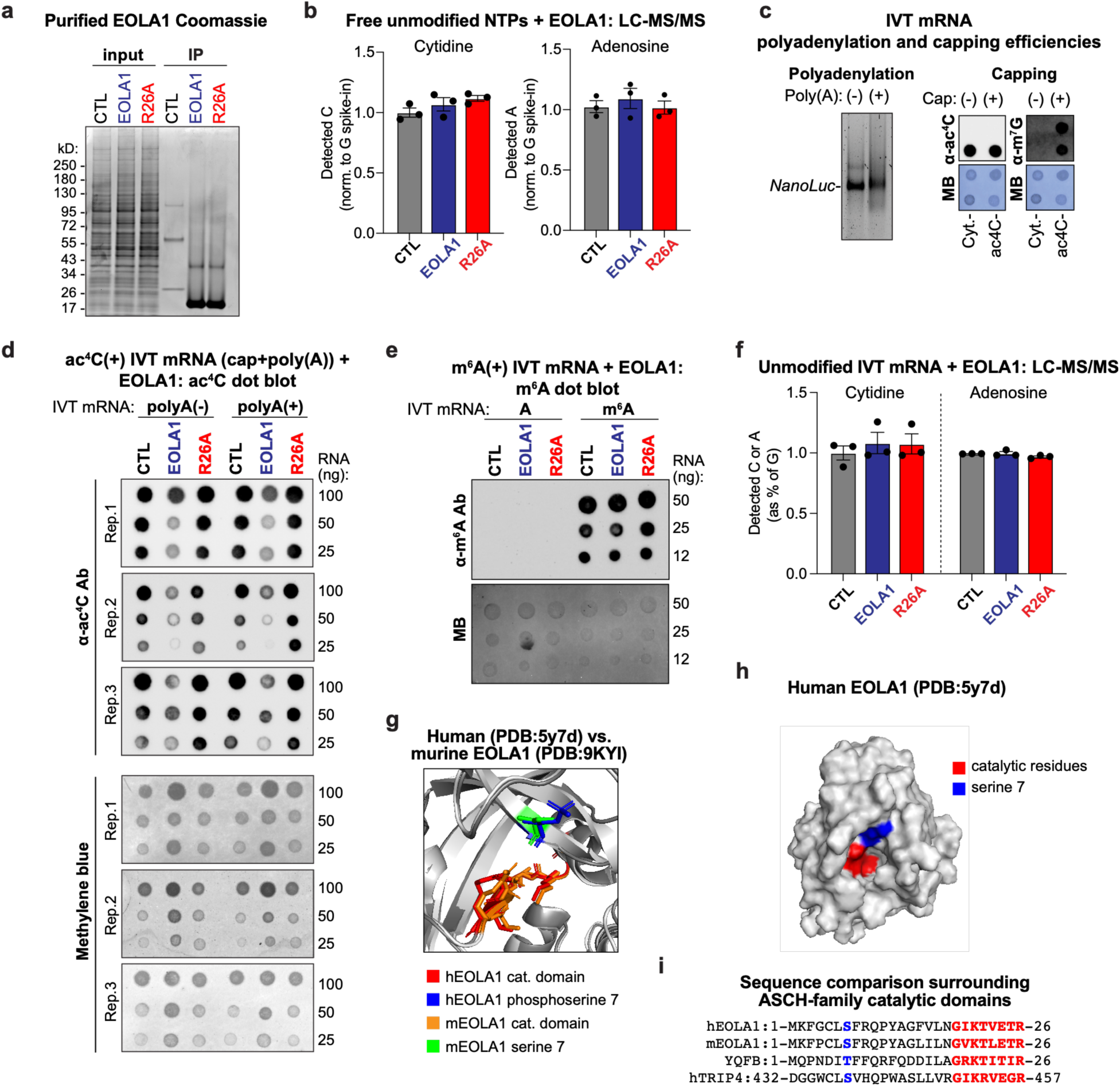
EOLA1 activity to ac4C in vitro. **(a)** Coomassie staining showing purity of EOLA1 following FLAG immunoprecipitation. **(b)** LC–MS/MS quantification of unmodified NTPs following ac^4^C treatment with wild-type EOLA1 or the R26A catalytic mutant. Data are mean ± SEM (n = 3); two-sided unpaired t-test; n.s. throughout. **(c)** Agarose gel analysis of in vitro transcribed *NanoLuc* mRNA. Polyadenylation was evaluated by a gel migration shift. Capping and acetylation were assessed through ac^4^C and m^7^G-dot-blot assay. **(d)** Dot-blot analysis of ac4C levels following EOLA1 treatment of *NanoLuc* mRNA with or without a poly(A) tail; methylene blue staining used as an RNA loading control. **(e)** Dot-blot analysis of RNA m^6^A levels following EOLA1 treatment; methylene blue staining used as an RNA loading control (representative of three experiments). **(f)** LC–MS/MS quantification of unmodified NTPs following m^6^A treatment with wildtype EOLA1 or the R26A catalytic mutant. Data are mean ± SEM (n = 3); two-sided unpaired t-test; n.s. throughout. **(g)** Structural representation comparing catalytic domains from human and murine EOLA1, showing phosphorylated Ser7. **(h)** Surface representation of the EOLA1 binding pocket highlighting phosphorylated Ser7. **(i)** Sequence alignment of the N-terminal ASCH domain showing conservation of Ser7; catalytic domain and Ser7 are indicated.

**Figure S6.**
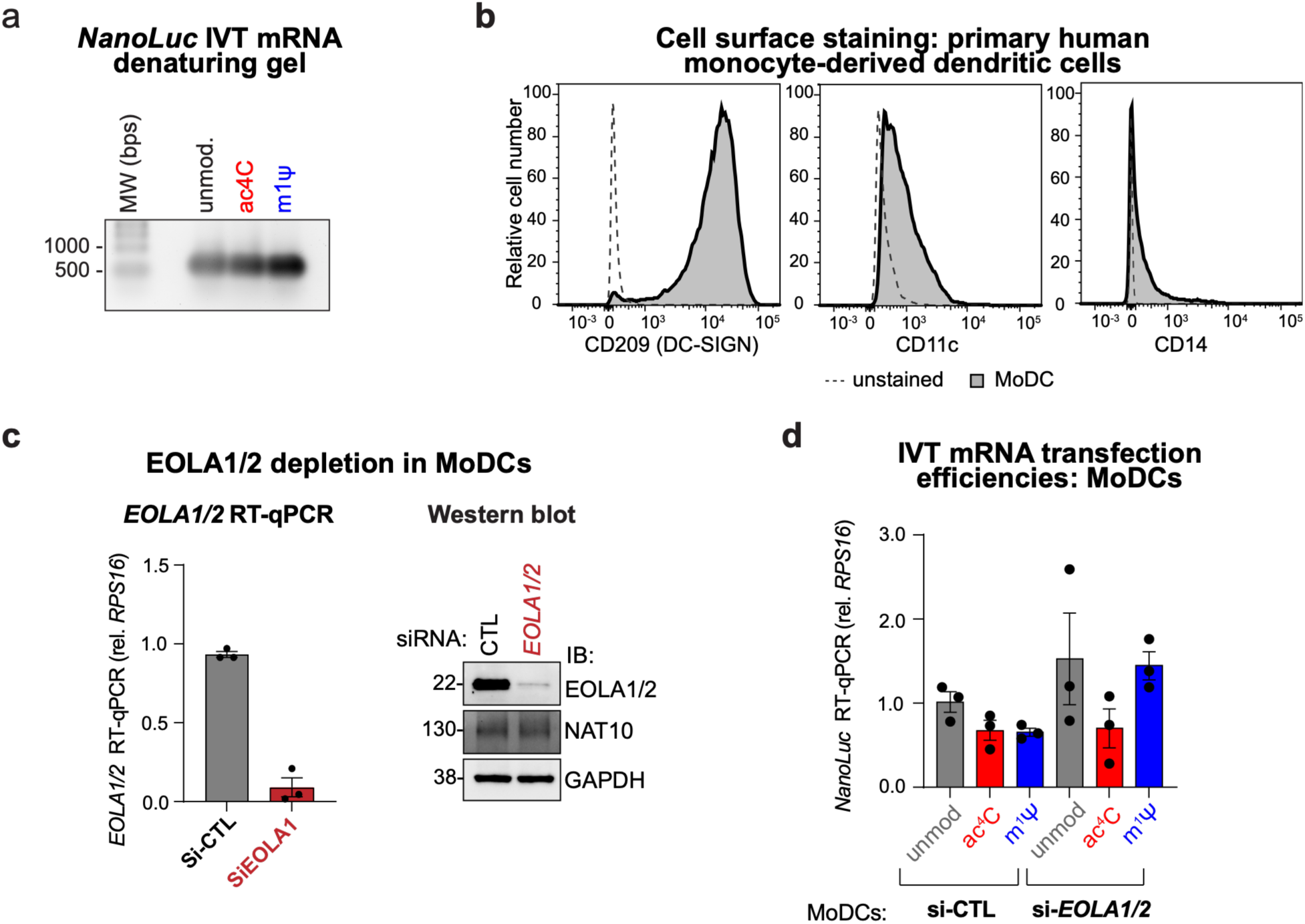
EOLA1/2 activity against synthetic mRNA in vivo. **(a)** Denaturing agarose gel electrophoresis of in vitro–transcribed *NanoLuc* mRNA. **(b)** Flow cytometry analysis of surface markers confirming differentiation of monocytes into monocyte-derived dendritic cells (MoDCs). **(c)** RT–qPCR and immunoblot validation of EOLA1/2 silencing in MoDCs (mean ± SEM., n = 3). **(d)** RT–qPCR assessment of *NanoLuc* transfection efficiency in MoDCs (mean ± SEM., n = 3).

